# Micro16S: Universal Phylogenetic 16S rRNA Gene Representations for Deep Learning of the Microbiome

**DOI:** 10.64898/2026.03.21.713432

**Authors:** Haig V. Bishop, Craig W. Herbold, Renwick C.J. Dobson, Olivia J. Ogilvie

## Abstract

Existing self-supervised microbiome models represent taxa as discrete, independent units restricted to fixed vocabularies, disregarding their evolutionary context. Here we present Micro16S, a deep learning approach that embeds 16S ribosomal RNA gene sequences into a continuous vector space according to phylogenetic relationships derived from the Genome Taxonomy Database. Using a combination of triplet and pair loss objectives, the model learns representations where spatial proximity reflects phylogenetic relatedness, while remaining largely invariant to the specific 16S rRNA region. Evaluations demonstrate taxonomically coherent clustering across most ranks and substantially improved region invariance compared to *k*-mer frequency baselines. A transformer pretrained on 50,418 unlabelled gut microbiome samples using these embeddings captured biologically meaningful community structure, though classical machine learning baselines outperformed Micro16S across six benchmark classification tasks, highlighting the limitations of the current system. These results establish the feasibility of phylogenetic embeddings for microbiome deep learning and identify mining algorithm design and class imbalance as primary targets for future improvement.

## 2 Introduction

The 16S ribosomal RNA (rRNA) gene, approximately 1,500 bp in length, is a ubiquitous taxonomic marker found throughout the tree of life including all prokaryotes, a group that encompasses both bacteria and archaea^1–3^. Its structure consists of nine hypervariable regions (V1 to V9) separated by highly conserved sequences. These conserved regions enable the design of universal primers that target particular 16S rRNA regions with broad effectiveness across prokaryotes. Meanwhile, the hypervariable regions provide the sequence diversity necessary to act as unique molecular identifiers across different organisms. In this way, microbiologists can use a single set of primers to amplify and sequence a target region, such as the hypervariable V3–V4 region, and identify the organisms based on the variants present. These unique structural features alongside the broad adoption by the field have resulted in 16S rRNA gene sequencing becoming the primary method for characterising microbial communities^3,4^.

Computationally, the first step to identify organisms in a sample from amplicon data involves processing raw sequencing data to infer the 16S rRNA gene variants present. A popular tool for this is DADA2, which identifies amplicon sequence variants (ASVs) and calculates their relative abundances^5^. To assign taxonomic identities to these ASVs, most pipelines employ a naïve Bayesian classifier, a method pioneered for this application by Wang et al.^6^ in the form of the Ribosomal Database Project (RDP) Classifier^6^. Within DADA2, this is reimplemented through the `assignTaxonomy` function. The algorithm operates by breaking sequences down into *k*-mers, which are short DNA sequences of a fixed length *k*, usually *k* = 8. The classifier then calculates the probability of a sequence belonging to a particular taxon based on how its *k*-mer frequencies compare to those in a reference database.

Several 16S reference databases exist, most notably Greengenes and SILVA^7,8^. These resources provide taxonomic classifications for 16S rRNA genes across a hierarchical system of seven ranks: domain, phylum, class, order, family, genus and species. However, these traditional taxonomies do not always reflect the true phylogenetic history of prokaryotes, largely driven by archaic nomenclature and biases introduced by older, less precise methods^9^. In 2018 the Genome Taxonomy Database (GTDB) addressed these issues by offering an alternative approach to prokaryotic taxonomy based on genome-wide information^10,11^. By using Relative Evolutionary Divergence (RED), the GTDB standardised taxonomic ranks so they represented consistent evolutionary depth across the entire tree of life. To support 16S-based research, the GTDB also provides a dedicated 16S rRNA gene reference database which aligns with its genome-derived classifications.

Collectively, these bioinformatic tools and 16S reference resources enable microbiologists to generate biologically relevant microbial community profiles. For examining microbial communities, the counts of each identified ASV can be used to produce a community composition. These profiles then fuel downstream statistical analysis or machine learning tasks^12,13^, enabling researchers to identify microbial community shifts for example, across different soil types^14^ or to predict clinical conditions, such as obesity, from a person’s gut microbiota^15^.

The simplest approach for applying machine learning to microbiome data involves arranging relative abundances into structured numerical lists called feature vectors to represent samples^16,17^. For 16S rRNA data, these features are typically ASVs or their assigned taxonomy at a specific rank, most often the genus. While functional, this approach has multiple limitations, stemming from the nuances of microbiome datasets. By restricting the model to a single taxonomic rank, the ability to differentiate lower-level taxa is lost, along with the broader evolutionary context provided by higher ranks. For instance, genus-level aggregation masks species-specific patterns, while treating taxa as independent features prevents models from generalising based on higher taxonomic levels and shared ancestry. Furthermore, the high-dimensionality of microbiome data often necessitates feature selection to avoid overfitting, which excludes rare but potentially significant organisms. Ultimately, this can leave models with a relatively narrow and imprecise view of the microbiome, limiting predictive performance.

To address these limitations, several phylogenetically informed machine learning methods have been proposed^18,19^, including Phylo-Spec and DeepBiome. Phylo-Spec integrates abundance data with phylogenetic relationships by guiding convolutions with a tree-traversal and incorporating phylogenetic distances^18^. On the other hand, DeepBiome employs a deep neural network architecture that directly mirrors the hierarchical structure of the phylogenetic tree^19^. While these approaches integrate taxon abundances with their phylogenetic context, they continue to treat taxa as single-dimensional, discrete units. In contrast, parallel developments have shifted toward representing the phylogenetic identity of taxa as points within a continuous, high-dimensional space^20,21^. DeepPhylo, for instance, uses principal component analysis to embed ASVs using distances derived from their phylogenetic tree^20^. This maps each ASV to a coordinate within a high-dimensional space where spatial proximity mirrors phylogenetic similarity. Woloszynek et al. (2019) adopted an NLP-inspired strategy, using word2vec skip-grams to embed ASVs based on their *k*-mer frequency profiles^21^. These high-dimensional vectors then allow downstream models to leverage dense sequence information and evolutionary context to inform classification tasks.

By integrating abundance data with phylogenetic relationships, these approaches avoid the limitations of relying on a single taxonomic rank, usually yielding higher performance on regression and classification tasks than simple abundance vectors. However, none of the above methods utilise the self-supervised learning paradigm, which involves pretraining foundation models that are later fine-tuned for specific tasks. This approach has transformed fields like computer vision and natural language processing (NLP), where models learn from vast amounts of unlabelled data, to develop highly robust and generalisable representations^22–25^. By learning the underlying structure of the data during pretraining, the model gains a sophisticated foundation that improves its ability to handle complex patterns in downstream tasks.

Recently, the field has shifted toward this self-supervised learning paradigm, with several pretrained transformer-based models for microbiome data emerging over the past year^26–29^. Pope et al.^26^ provided the first example, training an ELECTRA-style model on the 18,480 sample American Gut Project (AGP) dataset, using GloVe embeddings of the 26,726 ASVs^26^. While they acknowledged the limited scale and scope of the pretraining dataset, Pope et al.^26^ reported that the approach yielded robust prediction of irritable bowel disease that generalised across study populations. The largest scale example was the Microbial General Model (MGM), a transformer model which learned representations for 9,665 genera via next-token prediction pretraining on a massive corpus of 263,302 MGnify samples^28^. MGM outperformed state-of-the-art models like DeepPhylo^20^ on biome classification benchmarks and demonstrated impressive generalisation by maintaining high accuracy across geographic regions. Further, the BiomeGPT preprint expanded this approach to metagenomic profiles, successfully learning 2,168 species-level representations from 13,349 human gut samples^29^. Together, these models demonstrate the power of leveraging large-scale, unlabelled datasets to capture the intricate biological dependencies within microbial communities.

While self-supervised learning approaches such as those described above are powerful, leveraging vast amounts of microbiome data, they currently lack the phylogenetic information found in the approaches mentioned previously. A clear drawback of these models is that they are limited to learning representations for a finite set of taxa or ASVs. For instance, the GloVe embeddings used by Pope et al. (2025) are only effective for their specific dataset as they are strictly tied to the particular V4 16S rRNA gene region^26^. In the case of MGM and BiomeGPT^28,29^, because they use taxonomic assignments, they can be applied to new 16S or metagenomic datasets. However, they are still confined to a fixed number of genera and species, preventing them from incorporating the full breadth of a sample’s diversity. Further, during the learning process, these taxa are treated as independent units without incorporating their underlying phylogenetic relationships. By ignoring the hierarchical structure of the phylogenetic tree, these models fail to fully capture the evolutionary context that is fundamental to microbial biology.

We propose a novel strategy: phylogenetic embeddings that respect the hierarchical structure of the phylogenetic tree while simultaneously harnessing the self-supervised learning paradigm. Unlike existing methods, we take a more direct approach to embedding microorganisms within a sample by using original ASV nucleotide sequences as input rather than relying on taxonomic classifications or treating ASVs as discrete categories. By doing this we can natively support ASV sequences from any region of the 16S rRNA gene and avoid loss of information by collapsing data into independent units. Doing this requires a bespoke embedding model that can represent any 16S rRNA ASV as a vector where distance mirrors phylogenetic relatedness. By doing so, this approach avoids a reliance on discrete features derived from taxonomic assignments. Instead of training a model to learn fixed embeddings for a finite set of taxa, it enables the model to learn the importance of every clade and branch in the entire phylogenetic tree. This approach also aims to avoid the loss of species resolution due to genus-level aggregation, as seen in the MGM paper^28^, by preserving species-level precision within the embeddings whenever an ASV allows for it.

In this work, we evaluate the potential of a novel phylogenetic embeddings approach, implemented as Micro16S. This deep learning approach aims to improve upon existing methods by providing 16S rRNA gene embeddings that adhere to phylogenetic relationships while remaining invariant to the specific gene region. Leveraging the sequence data alongside genomically informed phylogenies and taxonomies from the GTDB, we trained a model to map 16S rRNA gene sequences into a compact 256-dimensional embedding space. We demonstrated that Micro16S embeddings approximately capture phylogenetic identity of input sequences across the taxonomic hierarchy, while remaining largely invariant to the specific gene region. Compared to the RDP Classifier, Micro16S demonstrated competitive taxonomic classification accuracy at the domain, phylum, and class levels, though performance was limited at lower ranks and in terms of macro accuracy. These embeddings were then used to pretrain a transformer model on 50,418 unlabelled gut microbiome samples from the Human Microbiome Compendium (HMC)^30^ to learn the structure of gut microbiomes. Finally, through fine-tuning on six benchmark tasks, we found that while Micro16S is capable of sample classification, its performance was generally exceeded by classical machine learning approaches. Ultimately, by developing a system that encodes 16S-derived sequences into universal numerical vectors representing phylogenetic relatedness, we suggest that further refinement of the Micro16S approach could enable superior performance in downstream tasks such as microbiome sample classification.

## 3 Results and Discussion

### 3.1 Dual-Loss Training Encodes Phylogenetic Identity into Vector Space

Using phylogenetic, taxonomic and sequence data from the Genome Taxonomy Database (GTDB), we trained two embedding models to generate vector representations for commonly amplified 16S rRNA gene regions. The two models, named the validation model and the application model, differ only in their training data composition. The validation model was trained and evaluated using a strict dataset split: an “excluded” set containing 1,002 sequences from 12 randomly selected families, with the remaining sequences split 80:20 into training (48,495 sequences) and testing (12,123 sequences) sets. Conversely, the application model was trained on the full dataset of 61,620 sequences to maximise its utility for downstream microbiome-level classification tasks. Model training was guided by a combination of triplet loss and pair loss.

Triplet loss, originally developed for computer vision tasks such as face recognition^31,32^, was adapted here to encode taxonomic relationships between 16S rRNA gene sequences (Fig. 1a). This mechanism operates on triplets of sequences at a particular taxonomic rank: the anchor sequence, the positive sequence, and the negative sequence. At the designated rank, the anchor sequence and the positive sequence are in the same taxon, whilst the negative is in a different taxon. The model is encouraged to position the embeddings so that the distance between the anchor and the negative is greater than the distance between the anchor and the positive plus some defined margin distance, thereby encoding the correct taxonomic relationship.

**Figure 1.**
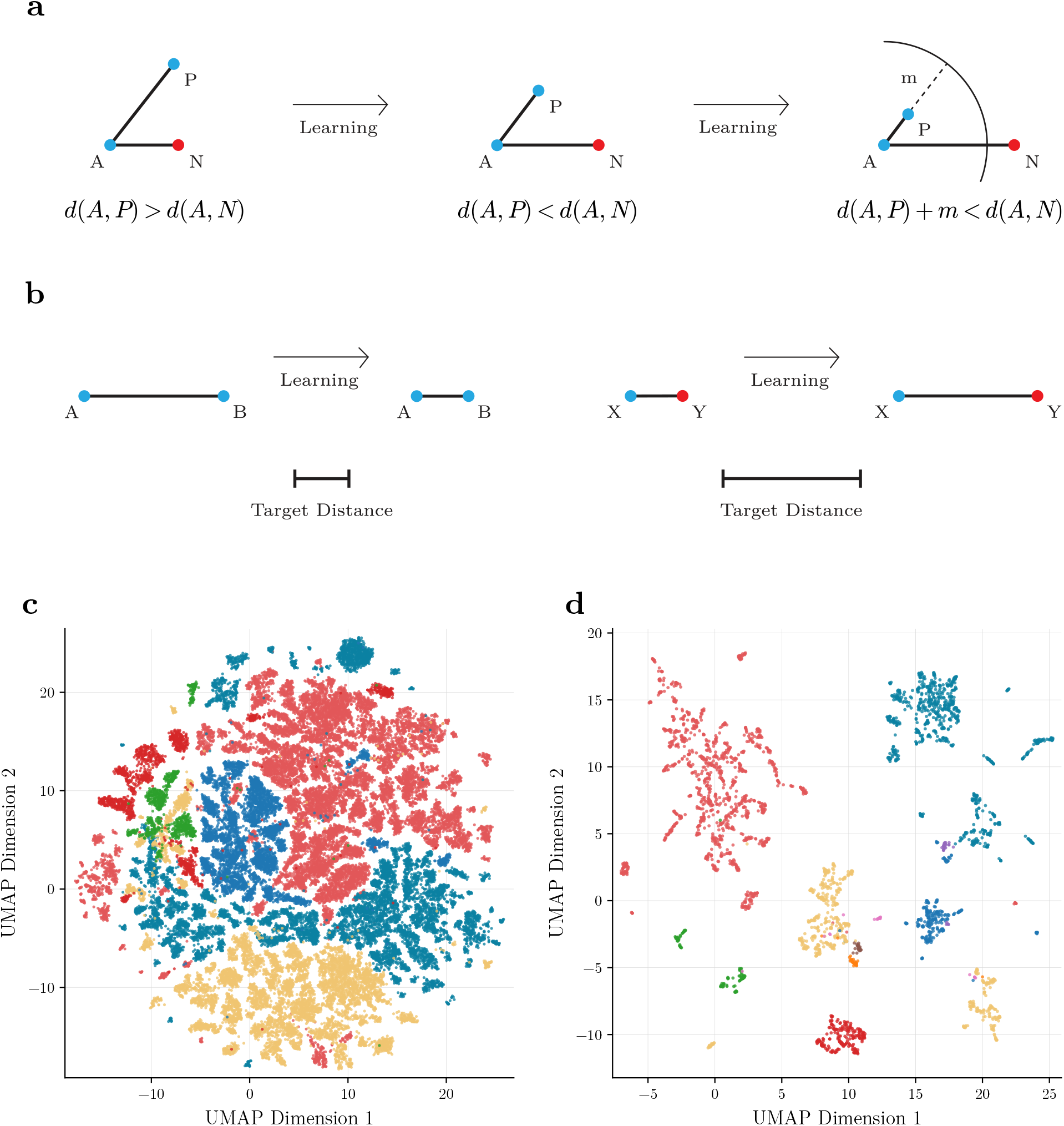
Training losses and UMAP visualisations for Micro16S embeddings. **a** Triplet loss mechanism used to separate embeddings by taxonomic relationships. Triplets consist of sequences called anchors (A), positives (P), and negatives (N). Learning is guided by the difference between the anchor-positive distance *d*(*A, P* ), the anchor-negative distance *d*(*A, N* ) and a margin *m*, where the model is encouraged to embed sequences such that *d*(*A, P* ) + *m < d*(*A, N* ). **b** Pair loss mechanism used to approximate pairwise phylogenetic relationships. Every pair of sequences is assigned a target distance derived from the GTDB phylogenetic tree. These target distances guide learning by encouraging the model to embed sequences either closer together or further apart to better represent the phylogenetic relationships. **c** UMAP projection of Micro16S application model embeddings for the six most abundant phyla, coloured by phylum. **d** UMAP projection of Micro16S application model embeddings for the ten most abundant families within Enterobacterales, coloured by family.

Complementing this, pair loss acts as a simple pairwise distance loss which regresses towards target distances between sequences (Fig. 1b). The model is encouraged to position sequences in the embedding space such that their pairwise distances match target distances derived from phylogenetic relationships provided by the GTDB. For instance, sequences differing at the phylum level are assigned larger target distances than those differing only at the genus level. Pair loss also extends to subsequences, where different regions of the same sequence are assigned a target distance of 0 to ensure the model recognises them as the same organism.

Together, triplet loss and pair loss steer the model to embed 16S rRNA gene regions so that phylogenetic identity is represented by spatial proximity, regardless of which specific sub-region of the 16S rRNA gene is provided as input. We visualised this using UMAP projections of the Micro16S application model embeddings from randomly selected training sequences. These include the six most abundant phyla (Fig. 1c) and the ten most abundant families within the order Enterobacterales (Fig. 1d). In both cases, embeddings of sequences from the same taxon tend to cluster together. In contrast, UMAPs of 7-mer representations of the same sequences are less coherently grouped according to taxonomy, particularly for the ten families in Enterobacterales (Appendix Fig. 1).

### 3.2 Embeddings Preserve Global Phylogenetic Structure and Reduce Region Bias

We found that Micro16S embeddings clustered broadly according to taxonomic identity across all ranks (Table 1). On the train and test sets, clustering performance was similar between the validation and application models, indicating that training the application model on the full dataset did not materially affect clustering performance. The largest differences in performance were observed across taxonomic ranks rather than between models or dataset splits. Phylum showed the weakest clustering, while performance was consistently higher for other ranks.

**Table 1.**
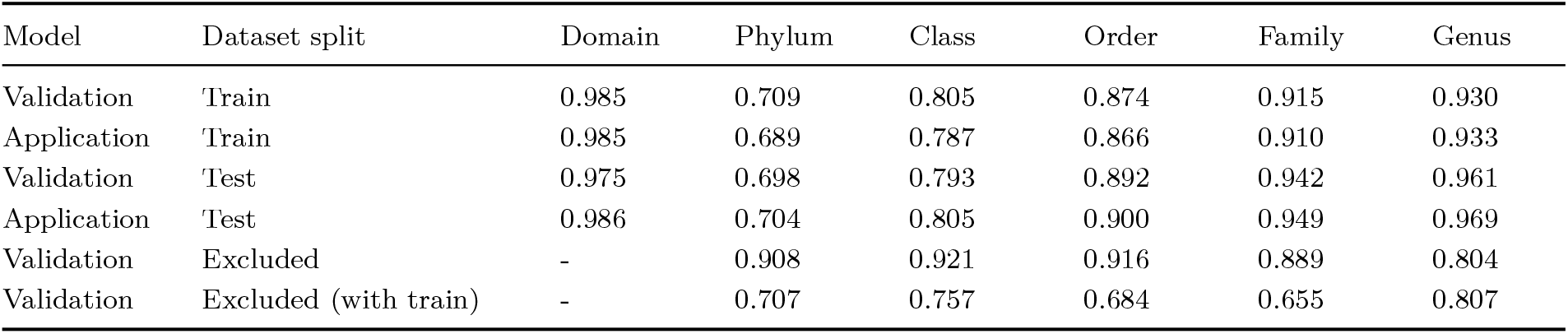
Clustering performance (V-measure) of Micro16S embeddings across taxonomic ranks for validation and application models on train, test, and excluded dataset splits.

We view this underperformance at the phylum level as a critical challenge for the success of Micro16S, as a robust top-level structure is essential to support the hierarchical substructures beneath it. We suspect this inferior performance at phylum rank stems from two primary factors. First, the sequence information in the 16S rRNA gene does not align well with the GTDB taxonomy. While phyla were traditionally defined by 16S rRNA gene sequences, the GTDB has updated these using genome-derived data^10^. Despite the training objectives of Micro16S leveraging the updated GTDB taxonomy, the model may still be heavily influenced by the inherent signal of the 16S rRNA gene. Second, there are a vast number of phyla with extreme class imbalance in the bacterial domain. The dataset used in this work contains 162 bacterial phyla with a median size of eight, and the ten largest phyla account for 89% of all bacterial sequences. We expect this skew, combined with the extreme sequence diversity at this rank, exposes the limitations of the current triplet and pair mining algorithm in providing an effective training signal.

When excluded families were clustered using embeddings from the validation model, performance was high at the phylum, class, and order ranks (Table 1). This suggested that the model retained the ability to infer broader phylogenetic structure for sequences from entirely unseen families, even when those taxa were absent during training. However, performance was lower at the family and genus ranks, demonstrating how generalisation was weaker at ranks where the model had not seen the taxa before. When the excluded sequences were clustered together with the training sequences, performance for the excluded set decreased at most ranks, particularly at phylum, class, order, and family. This indicates that although embeddings from unseen taxa still preserve some higher-level structure, that structure becomes less distinct when embedded alongside sequences spanning a broader taxonomic space.

The above evaluations all involve mixed random sampling across 16S rRNA gene regions, providing initial evidence of region robustness. To gauge the impact of using mixed regions, we performed additional clustering experiments using the validation model on the train and test sets, in which all sequences within a run were restricted to a single shared region. Compared with clustering under mixed-region sampling, this produced a slight increase in performance at all taxonomic ranks (Appendix Table 1), with the largest increase observed at genus level. This indicates that the Micro16S embeddings are largely, but not completely, invariant to the specific gene region used.

To quantify gene-region invariance more directly, we defined subsequence congruency (SSC), a measure of how similar embeddings from different regions of the same 16S rRNA gene are relative to embeddings from different 16S rRNA genes. Values approaching 1 indicate near-perfect congruency. On the train and test sets, for both validation and application models, SSC values were similar and high (Fig. 2). By comparison, 7-mer frequency vectors yielded substantially lower scores, demonstrating that Micro16S embeddings are more robust to region-specific signal. For the excluded families, SSC for the application model remained similar to that of the other dataset splits, whereas the validation model’s performance fell noticeably (Fig. 2). This reduction is consistent with the excluded families being entirely absent from validation-model training, limiting the model’s ability to place unobserved regions consistently in the embedding space.

**Figure 2.**
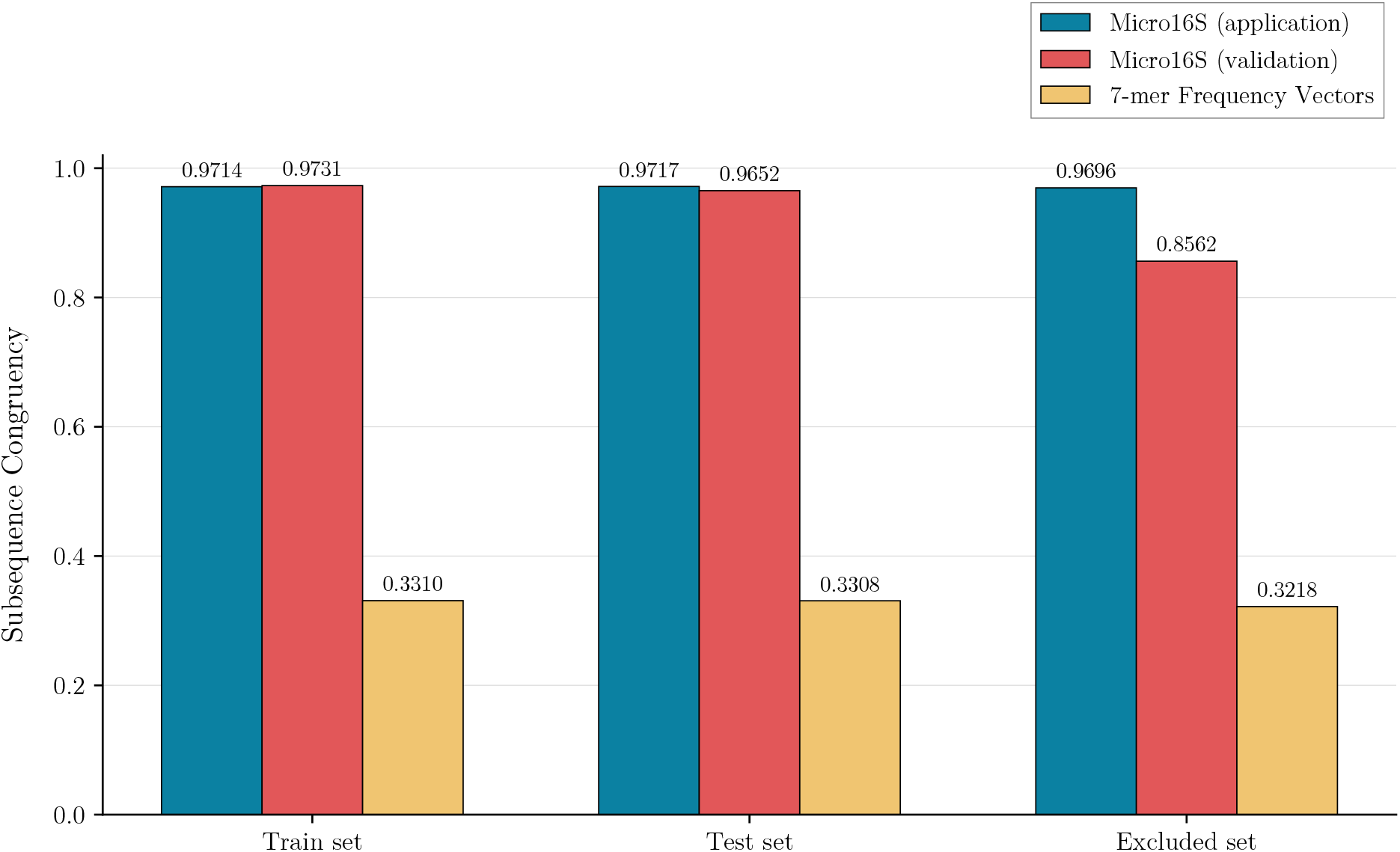
Subsequence congruency (SSC) scores of Micro16S embeddings and 7-mer baseline for train, test and excluded sets. SSC scores for train, test and excluded sets for all three methods with red bars showing Micro16S (validation model), blue bars showing Micro16S (application model), and yellow bars showing RDP.

### 3.3 Micro16S is Outperformed by Bayesian Method at Taxonomic Classification but Offers Reliable Confidence Scores

To assess Micro16S’s potential for usage as a taxonomic classifier, we compared its performance using a K-NN classification algorithm to the Ribosomal Database Project Naïve Bayesian Classifier (RDP), as implemented in DADA2. Classifying sequences in the test set using the training set as a reference, both the Micro16S validation and application models achieved accuracy scores lower than RDP at all ranks below domain, with the application model consistently outperforming the validation model (Fig. 3). The decrease in accuracy compared to RDP was modest at phylum-level, while the gap was far more pronounced at the lower ranks, particularly genus and species. Using macro accuracy to assess performance, which averages the accuracy across all taxonomic groups independently of their size, reveals an even more dramatic gap in performance of Micro16S compared to RDP. This suggests that Micro16S struggles to support classification of less common taxa despite the K-NN algorithm incorporating an inverse taxon-size weighting to votes (Appendix Fig. 2a).

**Figure 3.**
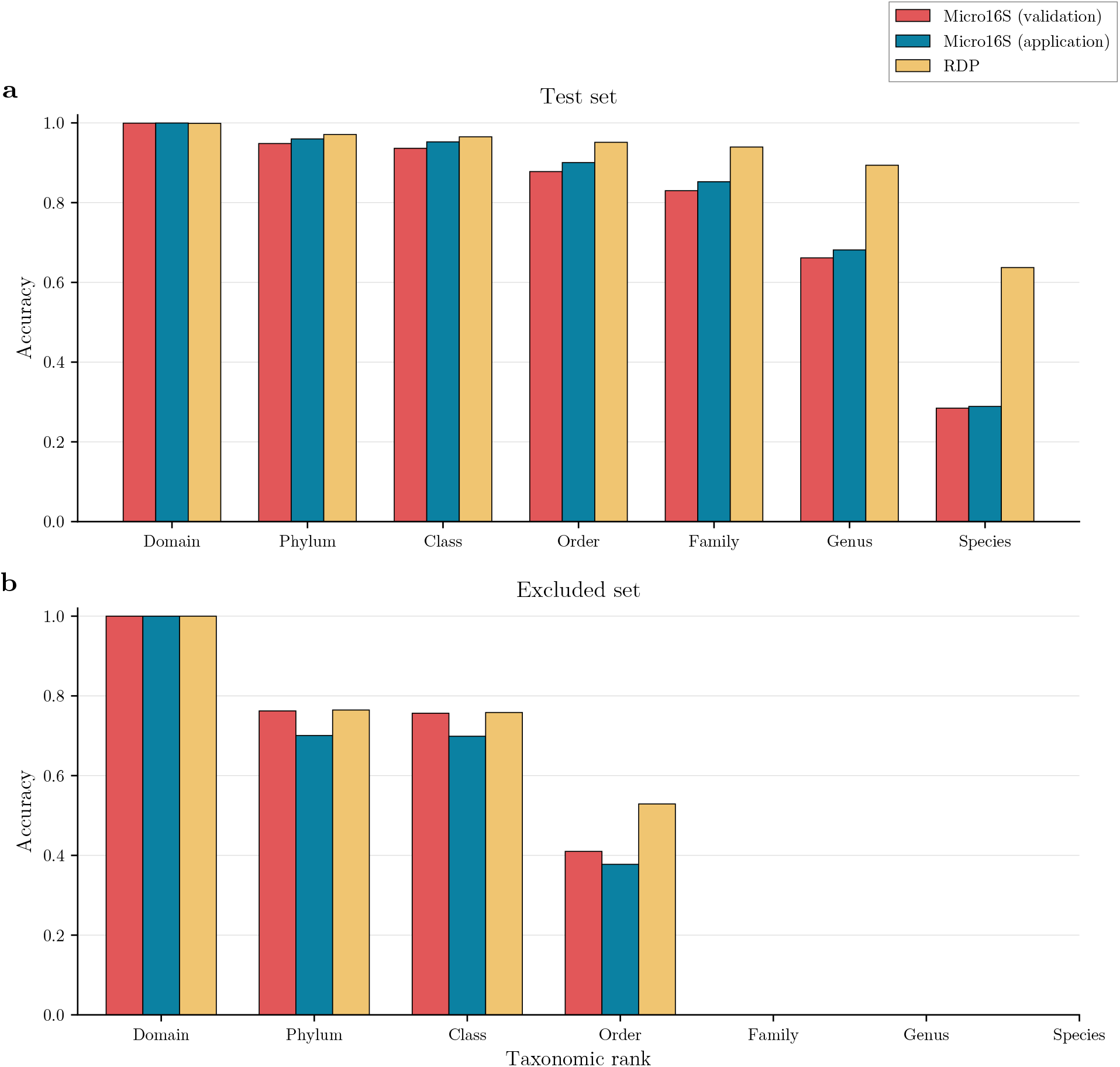
Accuracy scores for Micro16S and RDP across taxonomic ranks on the test and excluded sets. **a** Test set performance and b excluded set performance are shown as bar plots with taxonomic rank on the x-axis and pure accuracy on the y-axis. Red bars show Micro16S (validation model), blue bars show Micro16S (application model), and yellow bars show RDP.

We also evaluated Micro16S embeddings on taxonomic classification of the excluded set, where classification was only possible at ranks from domain to order, due to the exclusion of the families from the reference database. On these sequences, classification accuracy was much lower for all methods at ranks phylum, class and order (Fig. 3b), likely due to reliance on closely related sequences in the reference database. Curiously, on the excluded set the Micro16S validation model outperformed the application model. One possible explanation is that the application model had learned family-specific structure for these families during training. However, since those family-specific neighbours were absent from the classification reference set, the excluded sequences may have been embedded into relatively isolated regions reducing K-NN performance. Comparing the Micro16S validation model to the RDP algorithm revealed that at phylum and class ranks, sequences were classified with very similar accuracies between methods. However, this competitiveness diminished at the order rank, where Micro16S performance fell significantly below that of RDP.

The Micro16S K-NN taxonomic classification algorithm also produced confidence scores for every classification at every rank. By comparing confidence scores to observed accuracy, it was possible to see how well reported confidence aligns with the probability of correct classification (Fig. 4). This revealed that confidence scores aligned remarkably well with observed accuracy at all ranks. There was, however, some observed over-confidence at the lowest ranks, particularly species. With rank-wise correction, this shows promise that these confidence scores could act as reliable and intuitive indicators.

**Figure 4.**
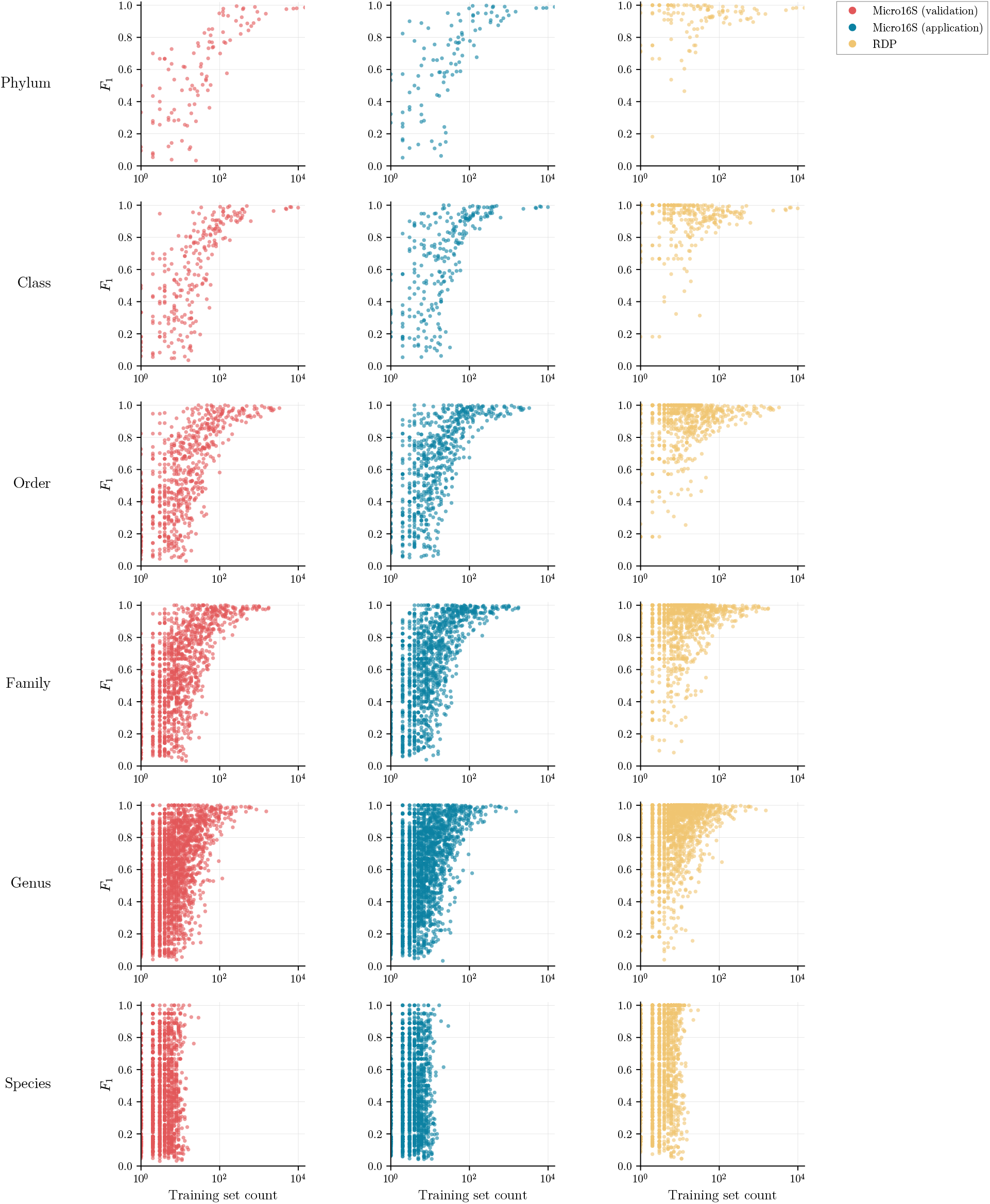
Confidence calibration plots for Micro16S models. One plot is shown per taxonomic rank from phylum to species, showing the relationship between confidence score and observed accuracy. Red lines show Micro16S (validation model), blue lines show Micro16S (application model). The black dotted lines show a perfect relationship between confidence and accuracy as reference.

To investigate how well Micro16S and RDP discriminate between taxonomic groups of different sizes, we compared taxon size to per-taxon *F*_1_ scores, which summarise how well each taxon is classified by combining two aspects: how often predictions for that taxon are correct, and how often sequences from that taxon are successfully detected. This revealed a clear pattern: as taxonomic group size increases, the lower bound of per-taxon *F*_1_ scores also increases, such that there are no large taxonomic groups with low *F*_1_ scores and the lowest *F*_1_ scores belong only to smaller taxonomic groups (Fig. 5). The observation that low per-taxon *F*_1_ scores were concentrated among small taxa, while large taxa rarely performed poorly, suggests that the primary bottleneck for Micro16S is data availability and training objective rather than just the algorithmic architecture. For large taxonomic groups, the high density of sequence examples in the training set allows models to define robust boundaries during training. Further, the triplet and pair mining algorithm is inherently biased towards selecting common taxonomic groups, possibly neglecting the learning of rare taxa.

**Figure 5.**
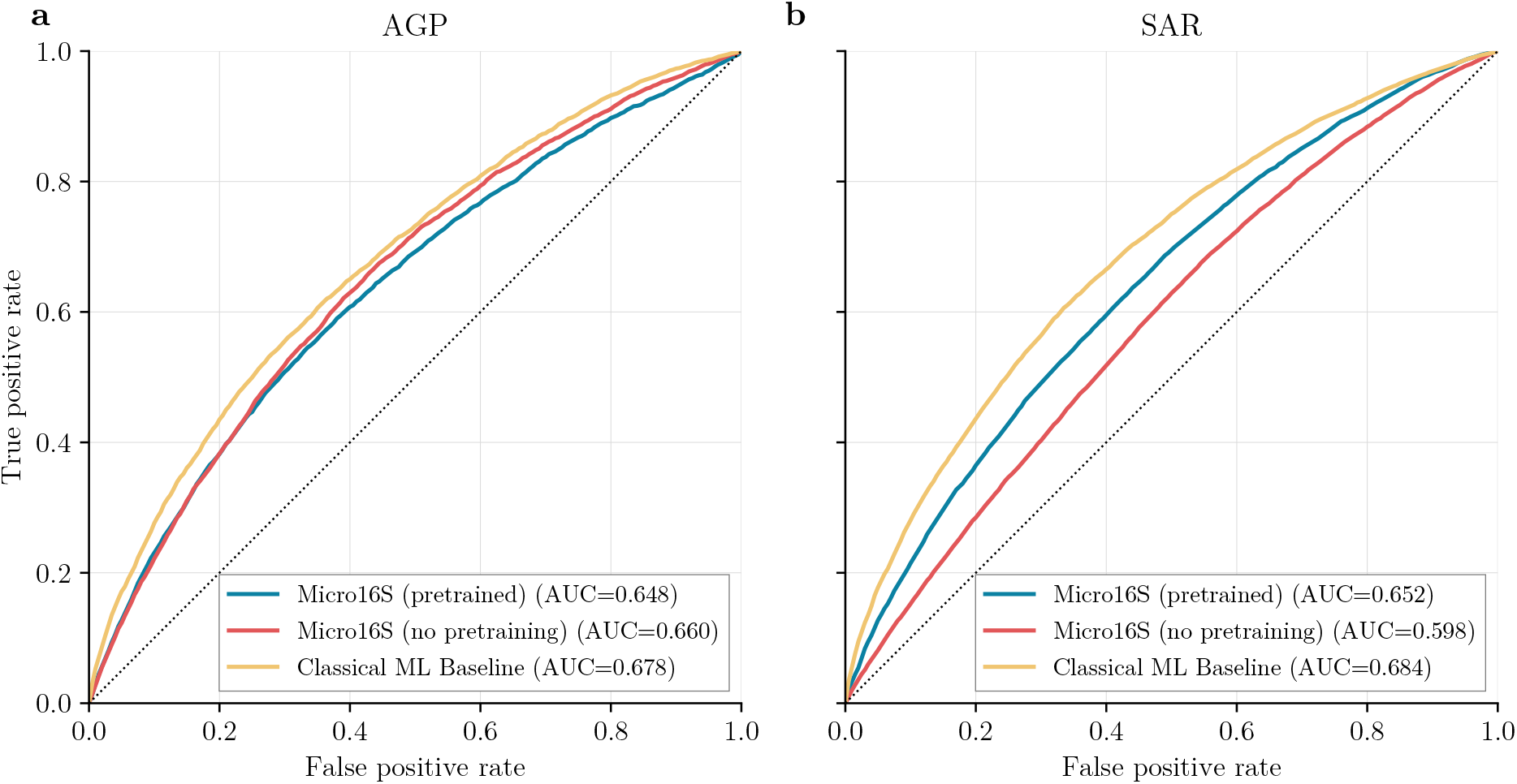
Per-taxon *F*_1_ scores by taxon size across taxonomic ranks and methods. Plots are shown for each rank (rows) and each method (columns), with the training-set taxon size (log scale) of taxa plotted against their per-taxon *F*_1_ score. Red denotes Micro16S (validation model), blue denotes Micro16S (application model), and yellow denotes RDP.

### 3.4 Micro16S-Based Pretrained Transformers Capture Demographic Signals

Using the Micro16S application model, we pretrained the microbiome transformer on 50,418 unlabelled gut microbiome samples across 149 studies from the Human Microbiome Compendium (HMC)^30^. This form of self-supervised learning involved training the model to reconstruct randomly masked ASVs and their relative abundances based on the surrounding community context, thereby encouraging the model to learn the co-occurrence patterns and compositional structure of the human gut microbiome. Both training and validation losses converged steadily, with the validation loss closely tracking the training loss, indicating that the model captured generalisable microbiome structure. This pretrained model was then finetuned on multiple classification benchmarks and compared to classical machine learning baselines. For comparison, we also finetuned microbiome transformers that had not undergone pretraining.

First, using K-fold cross-validation we evaluated these methods on gut microbiome sample classification of obesity using the American Gut Project (AGP)^33^ (*N* = 6, 108) dataset, and sex using the Sex-Age-Region (SAR)^34^ (*N* = 6, 914) dataset. We found that all three methods performed very similarly on AGP, with the classical machine learning baseline achieving a mean AUC of 0.678, while the pretrained and non-pretrained variants of the Micro16S model achieved 0.648 and 0.660, respectively (Fig. 6a). The impact of pretraining was not apparent for the AGP benchmark, however it did provide an increase in performance for the SAR benchmark, achieving a mean AUC of 0.652 compared to 0.598 for the non-pretrained version (Fig. 6b). Despite this pretraining performance gain, the classical machine learning baseline remained the top-performing method for the SAR dataset with a mean AUC of 0.684.

**Figure 6.**
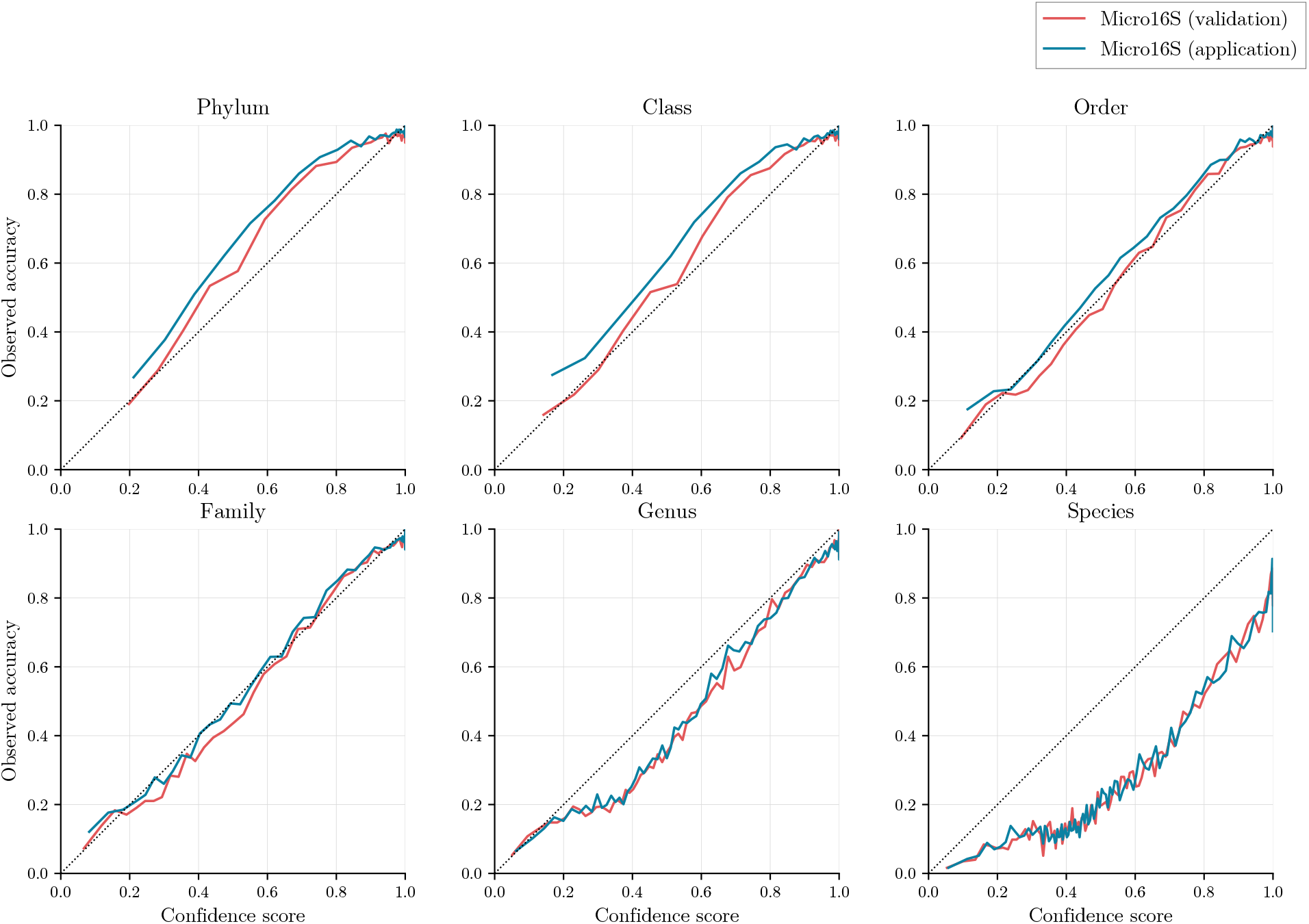
Microbiome sample classification performance on AGP and SAR benchmarks. **a** AGP ROC AUC plot for three methods: the pretrained Micro16S transformer (blue), the Micro16S transformer without pretraining (red), and the classical machine learning baseline (yellow). **b** SAR ROC AUC plot for the same three methods using the same colour scheme.

Overall, these demographic classification benchmarks showed that classical machine learning methods performed better than the current instantiation of the Micro16S-based method on microbiome-based demographic classification, and pretraining of the Micro16S-based transformer can provide enhanced performance in specific classification contexts.

### 3.5 Cross-Cohort Prediction of Celiac Disease Remains Limited for Micro16S-Based Transformer

To evaluate sample classification performance for predicting celiac disease within and between small heterogeneous cohorts, we utilised the data from the Celiac Microbiome Repository (CMR). These benchmarks represent a critical test of real-world applicability of microbiome-based machine learning methods to tasks where there is a limited supply of data from heterogeneous cohorts. Four separate classification tasks were constructed, each involving the combination of multiple celiac disease microbiome studies. All four tasks involved predicting celiac disease, with differences only being body site and disease stage: stool prospective, stool active, stool treated and duodenum active.

Using K-fold cross-validation, the classical machine learning baseline performed better than the pretrained Micro16S model on all four celiac disease prediction tasks (Fig. 7a). The best performances were seen on the stool treated task where the classical machine learning and Micro16S methods achieved mean AUCs of 0.892 and 0.822, respectively. In contrast, the stool prospective task yielded the lowest performances, with mean AUCs of 0.602 for the baseline and 0.525 for Micro16S.

**Figure 7.**
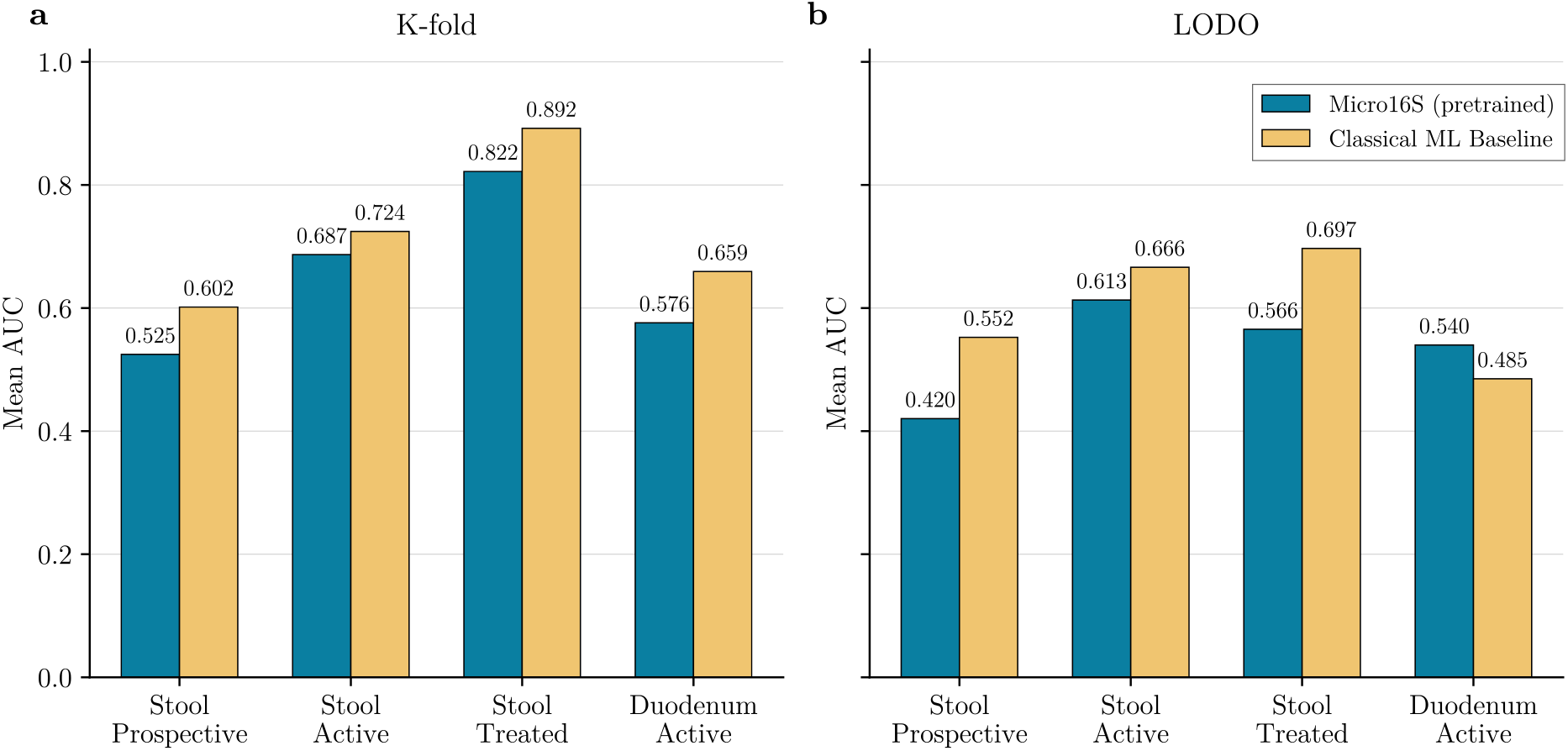
Microbiome classification performance on the CMR cross-cohort benchmark. **a** Mean AUCs over five K-fold cross-validation replicates across the four CMR-based celiac prediction tasks. The methods compared are the pretrained Micro16S transformer (blue) and classical machine learning baselines (yellow). **b** Mean AUCs over five LODO cross-validation replicates across the four CMR-based celiac prediction tasks, comparing the same methods using the same colour scheme.

Because these classification tasks each consisted of multiple studies, we were able to assess cross-cohort classification accuracy using leave-one-dataset-out (LODO) cross-validation, which better mimics real-world application of the model. Except for the duodenum active task, the Micro16S model consistently produced lower mean AUCs than the classical baseline (Fig. 7b). Overall, while the classical method showed modest success with LODO mean AUCs between 0.485 and 0.697, the Micro16S model struggled to generalise across cohorts, returning mean AUCs between 0.420 and 0.613.

### 3.6 Micro16S Establishes the Feasibility of Phylogenetic Embeddings

The central objective of Micro16S was to test whether a deep learning model could learn to embed 16S rRNA gene sequences into a vector space that reflects phylogenetic relatedness, while remaining invariant to the specific 16S rRNA region. The results presented here demonstrate that this core idea has validity and is measurable. The UMAP projections showed taxonomically coherent structure, clustering scores indicated that embeddings broadly captured taxonomic organisation across most ranks, and SSC scores substantially exceeded those of the *k*-mer frequency baseline. Crucially, when sequences from entirely excluded families were embedded by the model, their broader phylogenetic structure at the phylum, class, and order levels was present, indicating that the model has learned generalisable representations rather than simply memorising its training data. Though performance did not exceed classical methods, the ability of transformer models to learn demographic and disease-associated patterns from these embeddings further supports the view that they capture biologically meaningful information.

To the best of our knowledge, this is the first time a deep learning model has been trained to embed amplicon nucleotide sequence data into a vector space organised by genomically informed phylogenetic relationships. Previous embedding strategies for 16S rRNA genes have either derived distances from the ASV sequences themselves^20,21^ or have treated individual taxa or ASVs as discrete, independent units without any representation of their evolutionary relationships^26,28,29^. Instead, Micro16S takes a more direct approach of reading the raw nucleotide sequence and placing it in a space defined by the GTDB phylogenetic tree, built from genomic data rather than the 16S rRNA gene alone. A practical consequence of this design is that any 16S rRNA gene region can be projected into the same 256-dimensional space, allowing data generated from different amplicon protocols to be integrated without forcing a shared feature set. In the context of downstream classification, this also means that a model trained on these embeddings implicitly learns which branches and clades of the prokaryotic tree of life are relevant to a given prediction task, rather than independently learning the relevance of a fixed vocabulary of taxa.

Despite these encouraging results, the work exposed limitations of the current implementation. Phylum-level clustering was noticeably weaker than at other ranks, which we attribute to two compounding factors: a misalignment between the intrinsic sequence signal of the 16S rRNA gene and the genome-derived phyla defined by GTDB, and the extreme size imbalance of bacterial phyla in the training dataset. Classification accuracy at lower taxonomic ranks remained well below that of RDP, particularly for rare taxa, reflecting the difficulty of learning precise boundaries in parts of the tree where training examples are sparse. Although SSC scores were high overall, we suspect they may conceal region-specific bias at lower taxonomic ranks. Future work should utilise more targeted congruency metrics that focus specifically on relationships between sequences at lower ranks rather than averaging across randomly sampled sequences. The contribution of pretraining was also not fully resolved, with performance slightly improving on the SAR benchmark but marginally declining on AGP, leaving its effect uncertain. Finally, when the Micro16S embeddings were used to provide microbiome features to a transformer, any inaccuracies in the embeddings were likely inherited by the downstream model. This may explain why in almost all benchmarks the Micro16S-based approach underperformed classical machine learning baselines.

### 3.7 The Performance Ceiling Reflects Fundamental Challenges in Mining and Data Imbalance

A notable observation across clustering, classification and subsequence congruency evaluations is that Micro16S never surpassed a certain performance ceiling even on the training set itself, indicating that there is some limiting factor. Because the embeddings do not achieve near-perfect evaluation results even on training data, any inaccuracies are necessarily inherited by the downstream microbiome transformer, likely explaining why Micro16S-based approaches were generally outperformed by classical machine learning baselines.

We expect the primary bottlenecks to be the mining algorithm and class imbalance. The triplet and pair mining algorithm must simultaneously manage pair and triplet hardness, appropriately balance coverage across taxonomic ranks, and search through a combinatorially vast space of candidates within a feasible computational budget. These competing demands make it difficult to provide a consistently effective training signal, particularly for rare taxa. This is compounded by the inherent imbalance of prokaryotic taxonomic group sizes. Just a handful of abundant lineages encompass the vast majority of sequences, and the mining algorithm is inevitably biased toward these common taxa despite the inverse taxon-size weighting applied during candidate selection. The per-taxon *F*_1_ scores support this interpretation, with low scores concentrated almost exclusively among small taxa.

It is also worth noting that the training dataset of 61,620 sequences was significantly cut-down from the original 1,053,664 sequences. Of the sequences removed due to absence from the GTDB phylogenetic trees, most could likely still be assigned a position based on species-level matches. For the sequences excluded due to region extraction failures, most were partial-length 16S rRNA genes that span usable hypervariable regions.

Beyond these practical limitations, the phylogenetic embeddings approach carries some inherent constraints. The information content of the 16S rRNA gene places a hard limit on phylogenetic resolution, and the 600 bp input window currently does not accommodate full-length gene inputs. The approach is restricted to amplicon sequencing and cannot be applied to shotgun metagenomic data, though with some small adaptation the approach could be applied to other marker gene regions such as the ITS region, and to alternative phylogenies and taxonomic references.

### 3.8 Conclusions and Future Directions for Phylogenetic Embeddings

Existing self-supervised microbiome models treat taxa as discrete, independent units confined to fixed vocabularies, disregarding their evolutionary context. Micro16S addressed this by embedding 16S rRNA gene sequences into a continuous embedding space according to GTDB-derived phylogenetic relationships. Our evaluations demonstrate that the core concept is viable, with embeddings producing taxonomically coherent structure. However, phylum-level clustering was weak, taxonomic classification fell well below RDP for rare taxa, and the Micro16S-based transformer was consistently outperformed by classical machine learning baselines across all six benchmarks.

These results nonetheless establish a foundation for future development. The primary bottleneck is the mining algorithm, which struggles to provide consistent training signal across the combinatorially vast candidate space, particularly for rare taxonomic groups. Future work should experiment with alternative mining strategies to alleviate class imbalance, while expanding the dataset beyond the 61,620 sequences. With these refinements, phylogenetic embeddings represent a promising direction for microbiome deep learning, encoding evolutionary context directly into feature representations and enabling downstream models to learn which branches of the prokaryotic tree of life are relevant to a given prediction task.

## 4 Methods

### 4.1 16S rRNA Gene Data

The 16S rRNA gene reference database and the archaeal and bacterial phylogenetic trees from the GTDB Release 226^11^ were used as the source data to train Micro16S. These provided 16S rRNA gene sequences, along with genome-derived taxonomic annotations and phylogenetic distance relationships. Of the 1,053,664 16S rRNA genes available in the GTDB release, 921,200 were removed for not being present in the GTDB phylogenetic trees, and a further 24,519 were discarded as exact sequence duplicates to minimise redundancy. This filtering yielded a set of 107,945 full-length 16S rRNA genes, each with a defined position within the GTDB phylogeny.

### 4.2 Hypervariable Region Extraction

The in-house `extract16s` tool was used to identify and extract commonly used 16S rRNA gene regions from the full-length 16S genes. First, to identify hypervariable regions most commonly targeted by real-world gut microbiome 16S rRNA gene sequencing studies, the 1,314,161 ASVs from 250 diverse human gut microbiome datasets were used as a representative survey of sequencing methodologies. These datasets comprised 249 studies from the Human Microbiome Compendium (HMC)^30^ alongside the American Gut Project (AGP) dataset^33^. All ASVs were aligned against both the archaeal and bacterial barrnap 16S rRNA gene Hidden Markov Models (HMMs) using HMMER. Two GTDB reference sequences were simultaneously aligned to the same HMMs to establish a mapping from HMM model coordinates to reference gene coordinates: a 1,538 bp *Escherichia coli* sequence (`RS_ GCF_003697165.2∼NZ_CP033092.2:458562-460099`) and a 1,476 bp *Methanobrevibacter smithii* sequence (`RS_GCF_000016525.1∼NC_009515.1:333483-334958`). ASV hits were retained according to HMMER significance and coverage thresholds (*E*-value ≤ 10^−4^ for archaeal hits; *E*-value ≤ 10^−5^ for bacterial hits; query coverage ≥ 0.6), and each passing ASV was assigned to the domain for which it received the higher bitscore. Per-dataset consensus regions were then derived by filtering low-bitscore ASVs and positional outliers before computing the median start and end coordinates on each reference gene. To minimise redundancy, regions with start and end coordinates on both reference genes lying within 25 bp of each other were merged into a single representative region, yielding 29 unique 16S rRNA gene regions each defined by coordinate ranges on both reference genes (Appendix Table 2). These 29 regions were all categorised into variations of V1–V2, V1–V3, V3, V3–V4, V4, V4–V5, and V5–V6 regions.

These consensus region coordinates were subsequently applied to the 107,945 full-length 16S rRNA genes using `extract16s`, which employed the same archaeal and bacterial barrnap HMMs to locate each region within every gene. Sequences were excluded if they did not span the entirety of all 29 regions, contained ambiguous bases within any extracted region, or if the length of any extracted region fell outside the permitted bounds. For each region, the minimum permitted length was set to 50 bp below the shorter of the two reference sequence region lengths (archaeal and bacterial), and the maximum to 50 bp above the longer. Of the 107,945 sequences, 61,620 passed all filters. For each of the 29 regions, subsequences were extracted according to their reference coordinates with 30 bp of flanking sequence retained as padding on each side, producing 29 region-specific FASTA files each containing 61,620 sequences. Summary statistics for extracted sequence lengths are provided in Appendix Table 2.

### 4.3 16S rRNA Gene Dataset Construction

The 61,620 sequences were partitioned into three non-overlapping splits: train, test, and excluded. To construct the excluded set, 12 families with between 64 and 128 sequences were randomly selected and all of their sequences were withheld from training and test sets. These families and their respective sizes are listed in Appendix Table 3. The remaining sequences were then randomly partitioned into train and test sets at an 80:20 ratio. The resulting split sizes were 48,495 training sequences, 12,123 test sequences, and 1,002 excluded sequences.

### 4.4 DNA Sequence Encoding and Augmentation

Each of the 29 extracted regions across all 61,620 sequences was encoded as a 3-bit masked nucleotide representation within a fixed-length window of 600 bp. The first bit served as a binary validity mask, where a value of 0 indicated a valid nucleotide position and a value of 1 indicated a masked or padding position. The remaining two bits encoded the nucleotide identity according to the mapping `A ↦ 00, C ↦ 01, G ↦ 10, and T ↦` 11. Encoding all regions across the full dataset produced a tensor of shape [29, 61620, 600, 3].

Three augmentations were applied to every sequence at each training step and during evaluation of Micro16S: truncation, shifting, and point mutation.

To enhance region invariance, each sequence was subjected to random truncation at both ends, covering the natural variation in start and end positions of real ASVs. For each sequence, independent truncation amounts *t*_*L*_ and *t*_*R*_ were drawn uniformly between 0 and 60 bp and applied to the left and right ends respectively, with the truncated positions replaced by padding tokens. Because all sequences were extracted with 30 bp of flanking padding on each side, this truncation effectively adds or subtracts 0-30 bp relative to the original region start and end positions.

To reinforce shift and positional invariance, a randomly selected 50% of sequences underwent random shifting. In this augmentation, the contiguous block of unmasked nucleotide positions was repositioned to a uniformly random offset within the total available padding slack remaining after truncation, so that the sequence content is preserved but its absolute position within the fixed-length window varies.

To simulate sequencing error and natural within-species sequence variation, each unmasked base was independently mutated with probability *p*_mut_ = 0.01. Upon selection for mutation, each base was replaced with one of the three alternative bases with equal probability 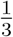, by randomly flipping one or both bits of the two-bit nucleotide encoding.

### 4.5 GTDB-Derived Phylogenetic Distances

The GTDB archaeal and bacterial phylogenetic trees were annotated with relative evolutionary divergence (RED) values using an in-house tool (`redvals`), following the RED definition introduced for GTDB rank normalisation by Parks et al.^10^. For each phylogeny (rooted as provided by GTDB), we set RED(*r*) = 0 for the root node *r*, and RED(*l*) = 1 for all leaf nodes (sequences) *l*. For all internal nodes *n* with parent *p*(*n*), RED was computed by linear interpolation along the lineage from *p*(*n*) to *n* according to the branch length to the parent, relative to the mean path length (sum of branch lengths) from the parent node to the extant descendant leaf nodes of *n*, denoted *u*(*p*(*n*), *n*).

Let *d*(*p*(*n*), *n*) denote the branch path length between *p*(*n*) and *n*, and let *L*(*n*) denote the set of leaf descendants of *n*. We define

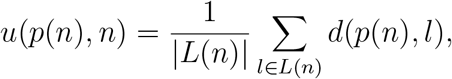

where *d*(*p*(*n*), *l*) is the path length from *p*(*n*) to leaf *l*. RED for *n* was then calculated as

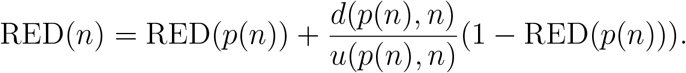

Pairwise RED distances between two leaf nodes (sequences) *i* and *j* were defined using the RED value of their most recent common ancestor (MRCA). The RED distance was

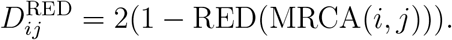

Pairwise RED distances were computed for all *N* = 61,620 sequences, yielding a symmetric distance matrix **D**^RED^ ∈ ℝ^*N ×N*^ . By construction, 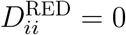because RED = 1 for leaves, and distances increase for pairs whose MRCA lies deeper in the tree, approaching 2 as the MRCA approaches the root.

To produce training targets that better facilitate learning of the taxonomic hierarchy in a cosine-similarity embedding space of 𝓁_2_-normalised vectors, RED distances were nonlinearly compressed to adjust resolution across taxonomic depths. Specifically, we first applied a global scaling factor *s* = 0.75 to all within-domain distances, then set cross-domain distances to a fixed constant *δ* = 1.6, and finally applied a *γ*-parameterised power transform:

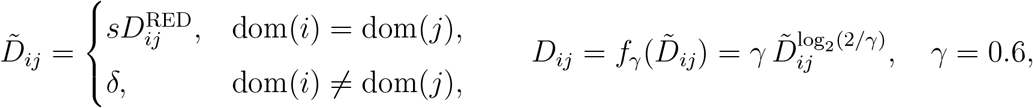

where dom(*·*)∈ {Archaea, Bacteria}denotes the GTDB domain label. This produced the final transformed distance matrix **D** used during training, which by construction, was constrained to the range [0, 2], matching the feasible interval for cosine-distances of 𝓁_2_-normalised embeddings (where 1− cos *θ* ∈ [0, 2]). In practice, this resulted in mean distances between taxa ranging between 1.3574 at domain rank, 0.0157 at species rank and 0 between subsequences. See Appendix Table 4 for summary statistics on all distances per rank.

### 4.6 Micro16S Architecture

#### Architecture Overview

Micro16S is a sequence-to-embedding neural network. Given a single 3-bit encoded nucleotide sequence of up to 600 bp, the model produces a single 256-dimensional,_𝓁2_-normalised embedding vector. The architecture is composed of five stages: (1) nucleotide embedding, (2) a convolutional stem, (3) a stack of transformer encoder layers each incorporating a depthwise convolutional sublayer, (4) attention pooling, and (5) an output projection head. The central design principle is the integration of local, motif-level feature extraction via depthwise convolution with global sequence-level context modelling via self-attention. To accommodate the variable-length sequences that arise from the use of different 16S rRNA gene amplicon regions, both operations are applied in a strictly padding-invariant manner throughout the network. In total, the model comprises 727,970 trainable parameters.

#### Nucleotide Embedding and Convolutional Stem

The 3-bit encoded input encodes a padding mask and a 2-bit nucleotide encoding. Positions are treated as a 5-token vocabulary *{*A, C, G, T, PAD*}*and mapped to learned embeddings with *d*_model_ = 96. The PAD embedding is fixed to zero and the mask is propagated so padded positions contribute no signal.

Embeddings are first processed by a mask-aware convolutional stem (1 *×* 1 expansion to 2*d*_model_ → Gated Linear Unit → (GLU) depthwise 1D convolution, *K* = **7**→**1** *×* 1 projection). Padding invariance is enforced by masking padded positions to zero throughout the stem and applying count-based renormalisation of the depthwise convolution outputs (scaling by *K/* max(*n*_*t*_, 1), where *n*_*t*_ is the number of valid tokens in the local kernel window). The stem is applied as a pre-norm residual block (**x** + *α·* conv(**x**)), with learnable *α* initialised to 0.1.

#### Transformer Encoder

The transformer encoder consists of a stack of four layers modelled on the Conformer architecture^35^. Each layer operates with model dimension *d*_model_ = 96, four attention heads (*d*_*h*_ = 24), and feedforward expansion dimension *d*_ff_ = 512. All layers follow a post-norm configuration, LayerNorm(**x** + Sublayer(**x**)), and contain three sublayers executed in sequence: (1) multi-head self-attention with Rotary Position Embeddings (RoPE), (2) a mask-aware depthwise convolutional sublayer, and (3) a position-wise feedforward network. In the self-attention sublayer, the input is projected to queries, keys, and values via a fused linear projection, reshaped into four heads, and attended using scaled dot-product attention. Position information is injected via RoPE^36^, which applies position-dependent rotations to query and key vectors such that each attention logit depends only on the relative offset between positions rather than on absolute indices, requiring no additional learned positional parameters. An additive mask sets logits at padded key positions to –∞, ensuring that padding contributes no signal after softmax normalisation. Head outputs are concatenated and projected back to *d*_model_.

The second sublayer is a mask-aware depthwise convolution, identical in design to the convolutional stem described above but with kernel size *K* = 15, not *K* = 7. By interleaving self-attention with depthwise convolution within each layer, the encoder jointly captures global sequence-level dependencies and local motif-level patterns, reflecting the core principle of the Conformer architecture.

The third sublayer is a position-wise feedforward network consisting of two linear projections with a GELU activation, expanding from *d*_model_ to *d*_ff_ = 512 and projecting back. As with the preceding sublayers, a residual connection followed by layer normalisation is applied.

#### Sequence Pooling and Output Projection

Next, the final encoder states for non-padded positions are pooled into a fixed-length representation using learned attention pooling, 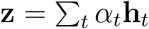, where *α*_*t*_ = softmax(*e*_*t*_ + *m*_*t*_) and *e*_*t*_ = **w**^T^ **h**_*t*_ + *b* (*m*_*t*_ = −∞ for padded positions and 0 otherwise). The pooled vector is passed through a two-layer projection head (96→ 96 → 256) with GELU to produce the embedding, which is then𝓁_2_-normalised to unit length, constraining all embeddings to the surface of a 256-dimensional unit hypersphere.

### 4.7 Micro16S Training Objectives

The model is trained with two complementary objectives: a pair loss that regresses embedding distances toward phylogenetic distance targets, and a triplet loss that enforces correct distance ordering across the taxonomic hierarchy. Both are formulated in terms of the cosine distance between 𝓁_2_-normalised embeddings,

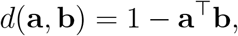

which yields values in [0, 2], matching the range of the transformed phylogenetic distance targets *D*_*ij*_.

Each pair and triplet used during training is associated with a taxonomic rank *r* from the GTDB hierarchy (domain, phylum, class, order, family, genus, species). For pairs, the rank is the level at which the two sequences first diverge: they share a common classification at all ranks above *r* but belong to distinct taxa at *r*. There is an additional dedicated “rank” of pairs called subsequences, comprising different 16S rRNA gene regions extracted from the same gene to enforce region-invariant embeddings. For triplets, the rank is defined by the anchor–negative relationship: the anchor and negative share classification at all ranks above *r* but differ at *r*, while the anchor and positive are required to share the same taxon at rank *r*.

Given a pair of embeddings **e**_*A*_ and **e**_*B*_ with target phylogenetic distance *D*_*AB*_, the pair loss is defined as the squared error between the predicted and target cosine distances, scaled inversely by the target distance and capped to bound outlier contributions:

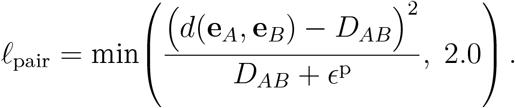

The denominator implements relative-error scaling: *ϵ*^p^ = 0.08 is a small constant that prevents gradient explosion for near-zero target distances. This formulation ensures that errors for closely related pairs are penalised proportionally more than those for distantly related pairs, directing optimisation effort toward finer-grained taxonomic distinctions. The cap at 2.0 limits the influence of any single outlier pair on the overall gradient.

Given a triplet of embeddings comprising an anchor **e**_*A*_, a positive **e**_*P*_ from the same taxon as the anchor at a given rank, and a negative **e**_*N*_ from a different taxon at that rank, the triplet loss enforces correct ordering of embedding distances. The per-triplet margin is set dynamically as the true phylogenetic distance gap, *m* = *D*_*AN*_ − *D*_*AP*_, and the loss is defined as

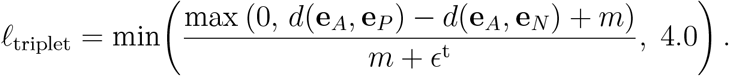

As with the pair loss, the hinge value is scaled inversely by the margin, with *ϵ*^t^ = 0.02 preventing division by zero. The inverse-margin weighting ensures that fine-grained triplets (e.g. at genus rank, where the positive–negative distance gap is small) carry proportionally larger gradients than coarse-grained triplets (e.g. at domain rank), concentrating learning on the most challenging taxonomic distinctions. The cap at 4.0 bounds the per-triplet contribution, stabilising training when margins are very small.

The total training loss is the unweighted sum of the mean pair and triplet losses over each batch:

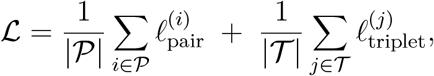

where 𝒫 and 𝒯 denote the sets of pairs and triplets in the batch, respectively. All linear and embedding layer weights are initialised with Xavier normal initialisation, all biases are initialised to zero, and the padding embedding vector is zeroed after initialisation.

### 4.8 Pair and Triplet Mining

#### Mining Overview

Both training objectives require selecting informative examples from the combinatorially vast space of possible pairs and triplets. We employ online mining, where at each training step the model’s current embeddings are used to identify the most useful examples. Each batch comprises 3,840 pairs mined across eight ranks (domain, phylum, class, order, family, genus, species and subsequence) and 7,680 triplets mined across six ranks (domain through genus). Both objectives follow a shared pipeline: candidate enumeration at each rank, subsampling to per-rank representative sets with taxon-size balancing, hardness computation and exponential moving average (EMA) based budget allocation, and percentile bucket sampling.

#### Sequence Selection and Distance Calculation

At each mining step, a random 20% of training sequences are subsampled, reducing the candidate pool and the computational cost of mining. Every subsampled sequence is then duplicated to provide material for subsequence pair mining. For each sequence and its duplicate, one of the 29 extracted regions is selected at random, with the constraint that each duplicate must receive a different region than its corresponding original. Augmentations (truncation, shifting, and point mutation, as described above) are applied independently to all sequences. The model then computes embeddings for the full set, from which a pairwise cosine distance matrix is calculated between all original sequence embeddings. Distances between each original and its corresponding duplicate are computed separately, yielding a cross-region cosine distance vector to be used for subsequence pair mining with a target distance of zero.

#### Per-Rank Budget Allocation

For both pairs and triplets, the total batch budget is distributed across the ranks in proportion to their current difficulty, so that ranks where the model struggles receive more training examples. Difficulty is tracked via an exponential moving average (EMA) of a per-rank hardness metric, updated after each mining step (see below) with smoothing factor *α* = 0.05.

For pair mining, the hardness metric at rank *r* is the sum of the 25th and 75th percentile squared relative errors, plus twice the mean squared relative error computed over the rank’s representative set:

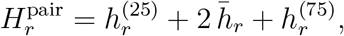

where *h* = *d*(**e**_*i*_, **e**_*j*_ − *D*_*i j*_) ^2^*/*(*D*_*i j*_ + *ϵ*^p^) as defined previously. For triplet mining, the hardness metric combines two proportions: triplets where the ordering is reversed (anchor closer to negative than positive), and triplets where the ordering is correct but the margin is violated (anchor too close to negative relative to positive):

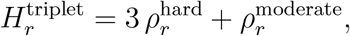

where 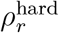 is the proportion of triplets at rank *r* for which *d*(**e**_*A*_, **e**_*P*_ ) ≥ *d*(**e**_*A*_, **e**_*N*_ ) (full violation), and 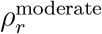is the proportion for which *d*(**e**_*A*_, **e**_*N*_ ) *< d*(**e**_*A*_, **e**_*P*_ ) + *m* (partial violation within the margin). The triple weighting on hard triplets directs additional budget toward ranks with outright ordering failures.

In both cases, the EMA-smoothed hardness values are normalised across the enabled ranks to yield budget proportions. Each rank is guaranteed a minimum share of 5% of the total budget to prevent any rank from being starved of training signal, while a 50% cap prevents any single rank from dominating the batch.

#### Pair Candidate Selection

At each rank *r*, all unique pairs of sequences *i* and *j* that share their taxonomic classification at all ranks above *r* but diverge at *r* were enumerated, with indices constrained to *i < j* to eliminate symmetric duplicates. Each rank’s candidate pool was then subsampled to a per-rank representative set of 40,000 candidates using weighted sampling without replacement. The weights counteract the combinatorial dominance of large taxa by downweighting pairs from abundant lineages:

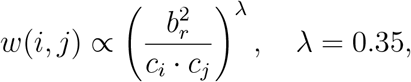

where *c*_*i*_ and *c*_*j*_ are the taxon sizes of sequences *i* and *j* at rank *r*, and *b*_*r*_ is the median taxon size at that rank across the training set. Subsequence pairs, drawn from the cross-region duplicate pool described above, are separately subsampled to up to 8,000 candidates.

For each candidate pair, the squared relative error *h* = *d*(**e**_*i*_, **e**_*j*_ − *D*_*ij*_) ^2^*/*(*D* + *ϵ*^p^) is computed as its hardness score, where *ϵ*^p^ is the same constant used in the pair loss. These scores drive both the per-rank EMA hardness update and the subsequent percentile bucket sampling.

#### Triplet Candidate Selection

At each rank *r*, all valid anchor-positive (AP) pairs were enumerated, where *A* and *P* share the same taxon at rank *r*, with the constraint *A < P* to eliminate symmetric duplicates. Each rank’s AP pool is then subsampled to a per-rank representative set of 40,000 candidates using weighted sampling without replacement. The weights follow the same taxon-size-balancing formula as pair mining, with *λ* = 0.8:

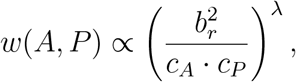

where *c*_*A*_ and *c*_*P*_ are the taxon sizes of the anchor and positive at rank *r*, and *b*_*r*_ is the median taxon size at that rank. Since the anchor and positive share the same taxon at rank *r*, these counts are actually equal.

Within each (anchor, rank) group, positives are then resampled using softmax-biased probabilities over the predicted anchor-positive embedding distance, *p*(*P* ) ∝ exp(*β*_*P*_ *·* .*d*(**e**_*A*_, **e**_*P*_ )) with *β*_*P*_ = 1.0, which upweights more distant (harder) positives while preserving the pool size. For each resulting AP pair, a negative is independently sampled from the set of valid candidates, defined as sequences that share classification with the anchor at rank *r* − 1 but differ at rank *r*. Negative selection also employs softmax-biased sampling, with probabilities *p*(*N* ) ∝ exp(– *β*_*N*_ *· d*(**e**_*A*_, **e**_*N*_ )) and *β*_*N*_ = 2.0, favouring closer (harder) negatives.

For each candidate triplet, the normalised violation is computed as

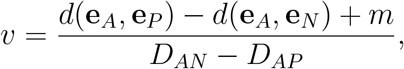

where the margin *m* = *D*_*AN*_ −*D*_*AP*_ as defined previously. Triplets with non-positive violation, which already satisfy the margin constraint and would produce zero loss, are filtered out to focus the training signal on informative examples. To prevent any rank from losing its candidate pool entirely, at least the size of the per-rank batch budget is retained after filtering. The normalised violations serve as the hardness scores for both the per-rank EMA update and the subsequent percentile bucket sampling.

#### Percentile Bucket Sampling

For both pairs and triplets, the final batch is assembled from each rank’s candidate pool via percentile bucket sampling. Candidates within each rank are sorted by their hardness score and partitioned into percentile-defined buckets, from which fixed proportions of the rank’s budget are drawn. Bucket boundaries are computed independently per rank, so that difficulty thresholds reflect the error distribution at each taxonomic depth to avoid being dominated by any single rank’s scale.

Pair mining uses six buckets defined at the 25th, 50th, 75th, 90th, 98th, and 100th hardness percentiles, with sampling proportions of 10%, 15%, 20%, 25%, 15%, and 15% respectively. This allocation emphasises moderate-to-hard errors while retaining a broad sample across the difficulty spectrum. Triplet mining uses four buckets at the 35th, 65th, 85th, and 100th percentiles, each sampled at an equal proportion of 25%. Because the hardest 15% of candidates receive the same share as the easiest 35%, this scheme biases sampling toward more challenging triplets. Although this is less aggressive than for pairs, it is counteracted by the removal of zero-loss triplets beforehand, meaning that buckets are computed only over non-zero loss examples. When a bucket contains fewer candidates than required, the shortfall is borrowed from adjacent easier buckets to maintain the target batch size.

### 4.9 Micro16S Optimization and Training

Both models were trained for 16,000 batches using the AdamW optimiser with a fixed learning rate of 2.5 *×* 10^−4^ and no learning-rate schedule. Each batch comprised 3,840 pairs and 7,680 triplets supplied by a single mining iteration. The two models were trained in parallel on the same system: a single NVIDIA GeForce RTX 4090 GPU with an Intel Core i7-14700F CPU and 128 GB of system memory. To accommodate GPU memory constraints, each batch was partitioned into 64 microbatches of 60 pairs and 120 triplets, with gradients accumulated across microbatches by scaling the loss by 1*/*64. Weight decay of 10^−3^ was applied selectively to embedding, linear, and convolutional layer weights, while biases, layer normalisation parameters, and the convolutional stem scalar gate were exempt. No dropout was used at any point in the network. Training employed CUDA automatic mixed-precision casting with bfloat16 for forward and backward passes, while the loss was computed in float32 for numerical stability.

### 4.10 Embedding Evaluations

All embedding evaluations were performed for both the validation and application models, with *k*-mer frequency vector baselines (*k* ∈ *{*5, 6, 7*}*) serving as a comparison. Only results for *k* = 7 are reported, because 7-mers consistently achieved the highest clustering V-measure and subsequence congruency scores across all dataset splits, while 8-mers were excluded due to memory constraints.

Two-dimensional UMAP projections^37^ were generated for application model embeddings and *k*-mer frequency vectors using training set sequences. Projections were computed with spectral initialisation, using cosine distance for Micro16S embeddings and squared Euclidean distance for *k*-mer vectors. Two visualisations were produced: 1) sequences from the six most abundant phyla, coloured by phylum (*n*_neighbors_ = 20, min_dist = 1.0). 2) sequences from the ten most abundant families within Enterobacterales, coloured by family (*n*_neighbors_ = 12, min_dist = 0.4). In both visualisations, each sequence was represented by one randomly selected region from the 29 available.

Clustering quality was assessed on the train, test, and excluded sets at all six ranks from domain to genus. The excluded set was clustered both independently and in combination with the training set, where in the latter case, scoring was calculated from only the excluded-set sequences. The number of clusters was set equal to the number of unique true taxa at the corresponding rank within the data being clustered, and cluster centres were initialised as the mean vectors of true-taxon members (label-centroid initialisation, *n*_init_ = 1). For Micro16S embeddings, spherical K-means was employed, with sequences and centroids 𝓁_2_-normalised at each iteration and cluster assignments determined by maximum cosine similarity. For *k*-mer frequency vectors, standard Euclidean K-means (Lloyd’s algorithm) was applied via scikit-learn. Both methods used max_iter = 200 and a convergence tolerance of 5 *×* 10^−4^. Clustering quality was quantified by V-measure. Two region-sampling strategies were employed: (1) a primary mixed-region analysis in which each sequence was assigned one of the 29 regions at random, with scores aggregated over 30 independent runs. (2) an analysis to probe region-invariance in which all sequences were assigned the same region per run across 29 runs (one per region, sampled without replacement), with scores aggregated.

To directly quantify the degree to which embeddings are invariant to the specific 16S rRNA gene region from which a sequence is derived, we introduce a metric termed subsequence congruency (SSC). SSC measures whether different region extractions of the same underlying 16S rRNA gene produce more similar representations than extractions from different genes. Formally, SSC is defined as

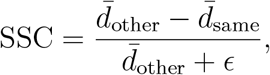

where *ϵ* = 10−^5^. The term 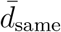 is the mean pairwise distance between the embeddings of all 29 regions extracted from the same gene, averaged across a set of sampled sequences, and quantifies how consistently a single gene maps to the same point in embedding space regardless of the region extracted. The term 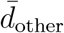 is the mean cross-sequence distance computed between all region-level embeddings from two disjoint sets of sampled sequences, providing a baseline measure of overall inter-sequence variability. Values approaching 1 indicate near-perfect region invariance, where the choice of 16S rRNA gene region has negligible influence on the resulting embedding relative to the differences between distinct sequences.

SSC was computed on the train, test, and excluded sets for both Micro16S models and for *k*-mer frequency vectors. Cosine distance was used for Micro16S embeddings and squared Euclidean distance for *k*-mer frequency vectors. For each combination of dataset split and representation, 20 independent runs were performed, each comprising 20 stochastic replicates. In each replicate, 1,000 distinct sequence indices were sampled without replacement: the first 500 were used to compute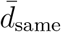 and the second 500 for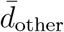. The run-level SSC was calculated as the mean across replicates, and the final reported value is the mean across the 20 runs.

### 4.11 Taxonomic Classification

To evaluate Micro16S’s potential as a taxonomic classifier, its performance was bench-marked against the naïve Bayesian classifier as implemented in DADA2’s `assignTaxonomy` function, here referred to as RDP^5,6^. The evaluation was performed for both the validation and application embedding models, on both the test and excluded dataset splits.

For each test or excluded sequence, 10 of the 29 extracted regions were randomly selected and the 30 bp flanking padding was truncated from each end to simulate realistic ASV boundaries, producing multiple region-specific query sequences per underlying 16S rRNA gene. For RDP, the training reference comprised the full-length 16S rRNA genes of all training set sequences. For Micro16S, because the 600 bp input window cannot accommodate full-length genes, each training gene was instead represented by a single randomly selected region.

RDP classification was performed via DADA2’s `assignTaxonomy` with default parameters (8-mer size, multithreading enabled, `minBoot =` 0), returning predicted taxonomy at all seven ranks along with bootstrap confidence values on a 0-100 scale.

Micro16S classification employed a weighted K-nearest neighbours (K-NN) algorithm operating on cosine similarities between𝓁_2_-normalised embeddings. Classification proceeded hierarchically from domain to species: at each successive rank, only training sequences whose predicted taxonomy matched at all higher ranks were retained as candidates, thereby enforcing taxonomic consistency across the hierarchy. The number of neighbours *K* was set per rank as [75, 50, 10, 10, 7, 5, 3] for domain through species respectively, reflecting the decreasing number of candidate training sequences at finer taxonomic depths. Each neighbour’s vote was weighted by two factors: (1) an inverse-square-root taxon-size weight, 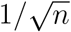, where *n* is the number of training sequences belonging to that neighbour’s taxon at the current rank, to mitigate the influence of class imbalance, and (2) an inverse squared cosine-distance weight, 1*/*(*d*^2^ + *ϵ*) with *ϵ* = 10^−6^, so that closer neighbours exerted greater influence on the prediction. The predicted taxon at each rank was the one receiving the highest sum of weighted votes. A confidence score was computed at each rank as the proportion of total weighted votes assigned to the winning taxon. To ensure that uncertainty at higher ranks propagated to finer ranks, the final reported confidence at each rank was defined as the cumulative product of per-rank confidence values from domain down to the rank in question.

For all classification metrics, sequences were excluded from evaluation at any rank where their true taxon was absent from the training set, since a classifier cannot be expected to identify taxa it has never observed. However, the same sequence could still contribute to metrics at higher ranks where its taxon was represented. Macro accuracy was computed as the mean of per-taxon accuracies at each rank. Per-taxon *F*_1_ scores were derived from per-taxon precision and recall, *F*_1_ = 2*PR/*(*P* + *R*), and were plotted against training-set taxon size. Confidence calibration was assessed by partitioning sequences into 100 percentile-based confidence bins and plotting the observed accuracy against the mean confidence within each bin.

### 4.12 Gut Microbiome Datasets

#### Human Microbiome Compendium (Pretraining)

The Human Microbiome Compendium (HMC) version 1.1.1^30^ was used as the unlabelled pretraining dataset. The HMC aggregates 16S rRNA amplicon sequencing data from hundreds of publicly deposited human-associated microbiome studies, providing harmonised ASV-level count tables and sample metadata. After quality filtering (minimum 1,000 total ASV reads and 50 unique ASVs per sample, minimum ASV length of 200 bp, exclusion of ASVs with ambiguous bases and datasets lacking a known amplicon region), 149 datasets comprising 50,418 samples across seven amplicon regions were retained. Datasets were split at the dataset level into training (*N* = 39, 733) and validation (*N* = 10, 685) sets at an approximate 80:20 ratio, ensuring no study appeared in both splits.

#### Sex-Age-Region Dataset (Classification)

The 16S rRNA amplicon dataset published by de la Cuesta-Zuluaga et al. (2019)^34^, hereafter referred to as the Sex-Age-Region (SAR) dataset, was used as a single-cohort classification benchmark. SAR comprises 7,009 stool samples from four countries (USA, UK, China, and Colombia) sequenced on the V4 region, with ASVs inferred by Deblur and trimmed to approximately 150 bp. Sample metadata includes sex, age, BMI, and country of residence. For the present work, SAR was used for binary sex classification (female vs. male). After preprocessing, 6,914 samples and 71,593 ASVs were retained, consisting of 3,812 females (55.13%) and 3,102 males (44.87%).

#### American Gut Project (Classification)

The American Gut Project (AGP)^33^ manuscript-package snapshot (1,250 rarefied release) was used as a classification benchmark. AGP is a large-scale citizen-science gut microbiome study providing V4 16S Deblur ASVs trimmed to 125 bp alongside extensive participant questionnaire metadata. For each sample, BMI was categorised into four groups: Under (BMI < 18.5), Normal (18.5 ≤ BMI < 25.0), Over (25.0 ≤ BMI < 30.0), and Obese (BMI ≥ 30.0). Samples were then filtered to retain only those in the Normal or Obese categories, with additional requirements of residence within one of the top seven countries by sample count in the dataset and a minimum of 50 non-zero ASVs per sample. The classification task was binary BMI category prediction (Normal vs. Obese). After filtering, 6,108 samples and 29,737 ASVs were retained, comprising 5,003 Normal (81.91%) and 1,105 Obese (18.09%) samples.

#### Celiac Microbiome Repository (Cross-Cohort Classification)

The Celiac Microbiome Repository (CMR) version 1.0 was used as a cross-cohort celiac disease classification benchmark. The following four 16S analysis groups were used: stool prospective (*N* = 120; 2 datasets), stool active (*N* = 158; 3 datasets), stool treated (*N* = 215; 4 datasets), and duodenum active (*N* = 118; 3 datasets). Each group aggregates multiple datasets generated using different sequencing protocols targeting the V3–V4/V4 region. Two versions of each group were prepared: truncated and untruncated. In the truncated version, ASV sequences from all constituent datasets were aligned and trimmed to their common overlapping V4 region, producing a harmonised ASV set suitable for classical machine learning models that require a shared feature space across cohorts. In the untruncated version, full-length ASV sequences were retained, relying on the region invariance of Micro16S embeddings to harmonise cross-cohort sequences with minimal information loss.

#### Profile Construction for Microbiome Transformer

For both pretraining (HMC) and finetuning (SAR, AGP, CMR), microbial profiles were constructed in a unified format. For each sample, raw ASV counts were normalised by total sum scaling, and the top *K* = 768 most abundant ASVs were selected and sorted in descending order of abundance. After truncation to the top *K*, abundances were re-normalised to sum to 1.0 over non-padded positions. Samples with fewer than *K* non-zero ASVs were zero-padded. Each retained ASV’s DNA sequence was embedded by the Micro16S application model, and the resulting vector was concatenated with the ASV’s normalised abundance, producing a per-sample profile matrix of shape [*K, d* + 1], where *d* is the embedding dimension. HMC profiles were split at the dataset level as described above. Finetuning datasets were constructed without predetermined splits, enabling flexible evaluation strategies (K-fold and LODO) to be applied at training time. For CMR, untruncated ASV sequences were used.

#### Profile Construction for Classical Machine Learning Baselines

For classical machine learning baselines, 2D abundance matrices were constructed. A shared ASV set was selected by retaining ASVs exceeding minimum average normalised abundance and sample prevalence thresholds. Raw counts were log_10_(*x* + pseudocount)- transformed without subsequent Z-score normalisation, and ASV filtering was applied after transformation. Each sample was represented as a vector of length equal to the number of selected ASVs, yielding a profile matrix of shape [*N*_samples_, *N*_ASVs_]. For CMR, classical machine learning models operated on truncated ASVs to enable cross-cohort comparison at the ASV level.

### 4.13 Microbiome Transformer Model Architecture

The microbiome transformer operates on microbial profiles represented as variable-length sets of ASV tokens. Each token comprises a 256-dimensional Micro16S embedding concatenated with the ASV’s normalised abundance. A linear input projection maps each token to the model’s hidden dimension (*d*_model_ = 256).

The projected tokens are processed by a transformer encoder comprising six layers, each with eight self-attention heads (*d*_*h*_ = 32) and a position-wise feedforward network with inner dimension 1,024 and GELU activation. Dropout of 0.1 is applied throughout, and layer normalisation follows each residual connection (post-norm). No positional encoding is used, as microbial profiles are unordered collections in which ASV position carries no biological meaning. Padded positions are excluded from attention computations via an additive mask.

Two task-specific output heads are attached to the encoder. For pretraining, an embedding reconstruction head (two-layer feedforward network: 256 → 1,024 → 256, GELU) projects each token back to the Micro16S embedding space, while an abundance reconstruction head (256 → 512 → 1) produces per-token logits normalised via a masked softmax over non-padded positions, yielding a valid compositional output summing to 1. For downstream classification, the per-token encoder outputs are aggregated into a fixed-length profile representation via learned attention pooling, in which a feedforward scoring network assigns per-token weights and the output is their weighted sum. A linear classification head then maps this pooled representation to task-specific logits. In total, the model comprises 7,079,426 trainable parameters.

### 4.14 Microbiome Transformer Pretraining

The transformer was pretrained on the HMC training set using masked autoencoding as a self-supervised objective, with the held-out HMC validation set used for model selection. At each training step, 15% of non-padded ASVs in each sample were randomly selected for masking. Both the Micro16S embedding and abundance of each selected ASV were replaced with fixed zero vectors, denying the model any direct information about the masked positions. ASV order within each profile was randomly shuffled at every step to reinforce the model’s permutation invariance.

#### Masked Token Matching and Loss

Because both the embedding and abundance are masked together and ASV order is meaningless, the model’s predicted reconstructions at masked positions have no positional correspondence to the ground-truth masked tokens. To resolve this assignment ambiguity, Sinkhorn optimal transport (OT) matching^38^ was used to align predicted masked tokens to their most probable ground-truth counterparts. For each sample, a pairwise cost matrix was computed over all masked positions, combining cosine distance for embeddings with absolute abundance error, each component normalised by its mean magnitude. A balanced transport plan was then computed via 25 Sinkhorn-Knopp iterations with entropic regularisation *ϵ* = 0.05. To further break symmetry between identical masked-token inputs and allow the model to produce distinct reconstructions for each, a random 16-dimensional code was sampled for each masked position at every step and concatenated to the token input prior to the input projection.

The total pretraining loss was a weighted sum of four terms:

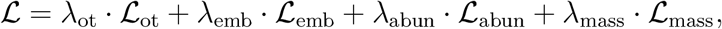

where ℒ_ot_ is the Sinkhorn OT cost (total transport cost under the plan, normalised by plan mass), ℒ_emb_ is the cosine similarity loss between soft-matched predicted and true embeddings, ℒ_abun_ is the mean squared error between soft-matched predicted and true abundances, and ℒ_mass_ is a mass consistency term penalising the absolute difference between total predicted and true abundance over masked positions. Loss weights were *λ*_ot_ = 0.1, *λ*_emb_ = 1.0, *λ*_abun_ = 200.0, and *λ*_mass_ = 1.0.

#### Optimisation

The model was trained for 16 epochs with the AdamW optimiser at an initial learning rate of 10^−4^ and weight decay of 10^−3^, using a batch size of 256 samples. A cosine annealing schedule with 1,500 linear warmup steps was applied over 15,000 total scheduler steps, decaying the learning rate to 10% of its peak value. Gradient𝓁_2_ norms were clipped to 1.0. Training used CUDA mixed-precision (bfloat16) on a single NVIDIA GeForce RTX 4090 GPU. The checkpoint at epoch 16 was selected for downstream finetuning.

### 4.15 Microbiome Classification Evaluation

#### Transformer Finetuning

The pretrained microbiome transformer was adapted for classification by replacing the pretraining reconstruction heads with the classification head described in the model architecture. The full model was finetuned end-to-end using the AdamW optimiser with learning rates 5 *×* 10^−5^ and 10^−4^ for the pretrained transformer body and randomly initialised classification head, respectively, alongside a weight decay of 10^−2^. The training objective was cross-entropy loss with inverse-frequency class weights to address class imbalance. Models were trained for 20 epochs without early stopping, with the final epoch’s model state used for evaluation. Batch sizes were 96 for the AGP and SAR datasets and 32 for the CMR benchmarks. ASV order within each profile was randomly shuffled at every training step. Training used CUDA mixed-precision (bfloat16) on a single NVIDIA GeForce RTX 4090 GPU. To isolate the contribution of pretraining, a non-pretrained control was evaluated on the AGP and SAR benchmarks, using an identical architecture initialised with random weights and the same training procedure.

#### Classical Machine Learning Baselines

Random Forest and XGBoost classifiers served as classical machine learning baselines, operating on the 2D log-transformed abundance matrices described above. For each benchmark, the best model type and hyperparameter configuration were jointly selected via exhaustive grid search with 5-fold stratified cross-validation, optimising macro-averaged AUC. The Random Forest search grid comprised `n_estimators` ∈ {100, 400}, `max_depth` ∈ {10, None}, `min_samples_leaf` ∈ {1,4},`max_samples` ∈ {0.7, 1.0}, and `max_features`∈{sqrt, log2}(32 configurations). The XGBoost search grid comprised n_estimators ∈ {200, 500}, `learning_rate` ∈ {0.05, 0.1}, `max_depth` ∈ {3, 6}, `min_child_weight` ∈ {1, 5},∈ `subsample` ∈ {0.8, 1.0}, and `gamma` ∈{0, 1}(64 configurations). Random undersampling of the majority class was applied independently to each training fold prior to model fitting. Across the six benchmarks, the selected model type was evenly split between Random Forest and XGBoost, with three wins each.

#### Evaluation Protocol

All classification benchmarks were evaluated using 5-fold stratified cross-validation repeated over five independent replicates, each with a distinct random seed for fold partitioning. For the four CMR classification tasks, leave-one-dataset-out (LODO) cross-validation was used to assess cross-cohort generalisation. In each iteration, all samples from a single study were held out for testing while the model was trained on the remaining studies, repeated for five replicates. For the Micro16S transformer, a fresh model was initialised from the pretrained (or randomly initialised) checkpoint at every fold or held-out group, ensuring independence across evaluations. For the machine learning baselines, the tuned model type and hyperparameters were used for all evaluation folds and groups. Class imbalance handling was applied exclusively to training data. The primary evaluation metric was ROC-AUC, and results are reported as mean AUC across replicates.

## 5 Data Availability

All source data used in this study are publicly available. The GTDB Release 226 16S rRNA gene reference database and phylogenetic trees were obtained from the Genome Taxonomy Database (https://gtdb.ecogenomic.org/)^11^. The Human Microbiome Compendium v1.1.1 is available via Zenodo (https://zenodo.org/records/15122187)^30^. The American Gut Project manuscript-package 1,250 rarefied release was retrieved from https://ftp.microbio.me/AmericanGut/manuscript-package/1250/^33^. The Sex-Age-Region dataset is available at https://github.com/jacodela/microbio_aDiv^34^. The Celiac Microbiome Repository is available at https://github.com/CeliacMicrobiomeRepo/celiac-repository.

## 6 Code Availability

The Micro16S embedding model source code and both trained model weights (validation and application) are available at https://github.com/HaigBishop/micro16s. The microbiome transformer pretraining and finetuning code, along with the pretrained model weights, are available at https://github.com/HaigBishop/micro16s-transformer. The tools `redvals` (RED value computation) and `extract16s` (16S region extraction) are available at https://github.com/HaigBishop/redvals and https://github.com/HaigBishop/extract16s, respectively.

## Appendix

**Figure 1.**
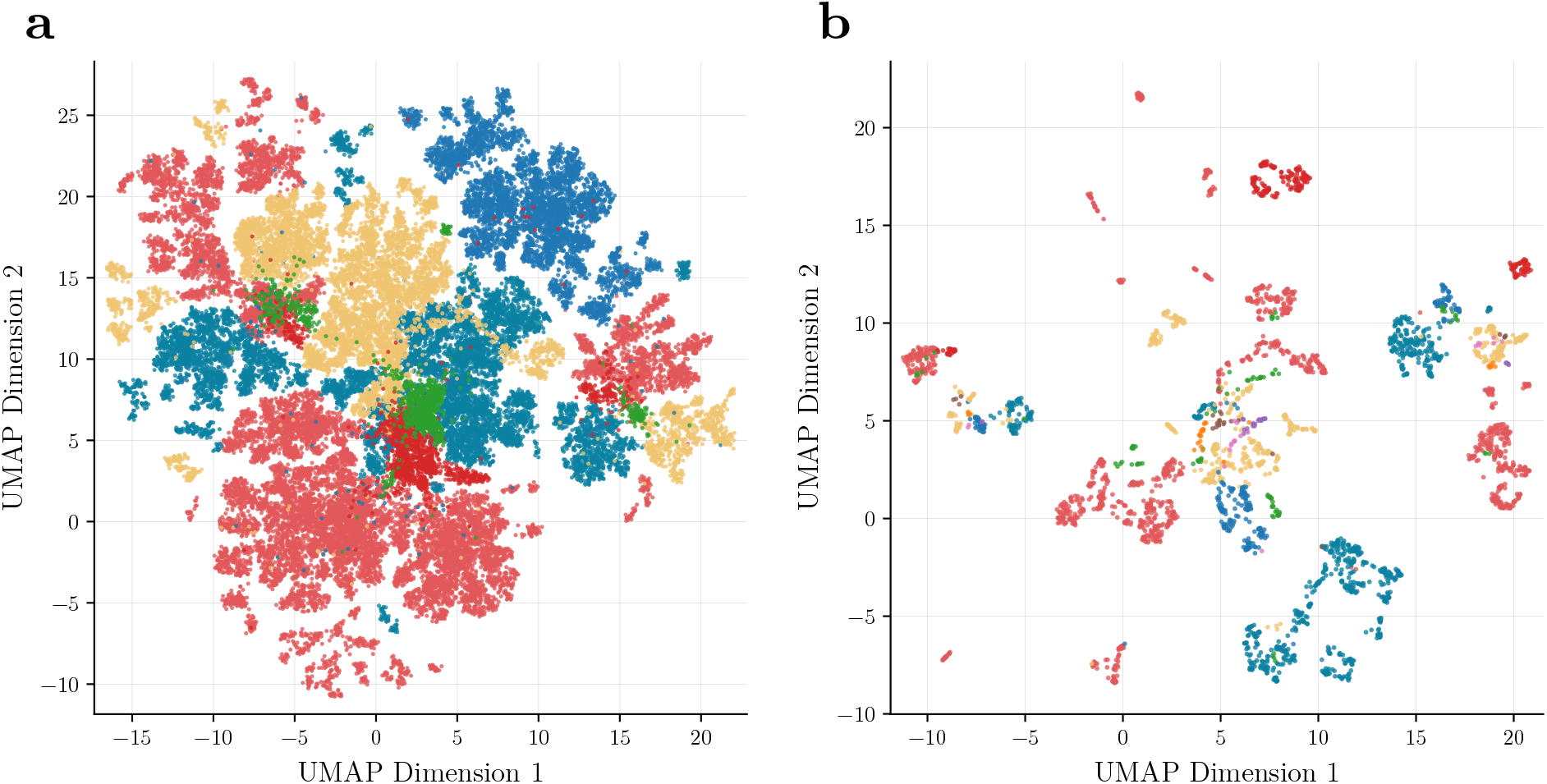
UMAP visualisations of 7-mer frequency representations. **a** UMAP projection of 7-mer frequency vectors for the six most abundant phyla, coloured by phylum. **b** UMAP projection of 7-mer frequency vectors for the ten most abundant families within Enterobacterales, coloured by family.

**Figure 2.**
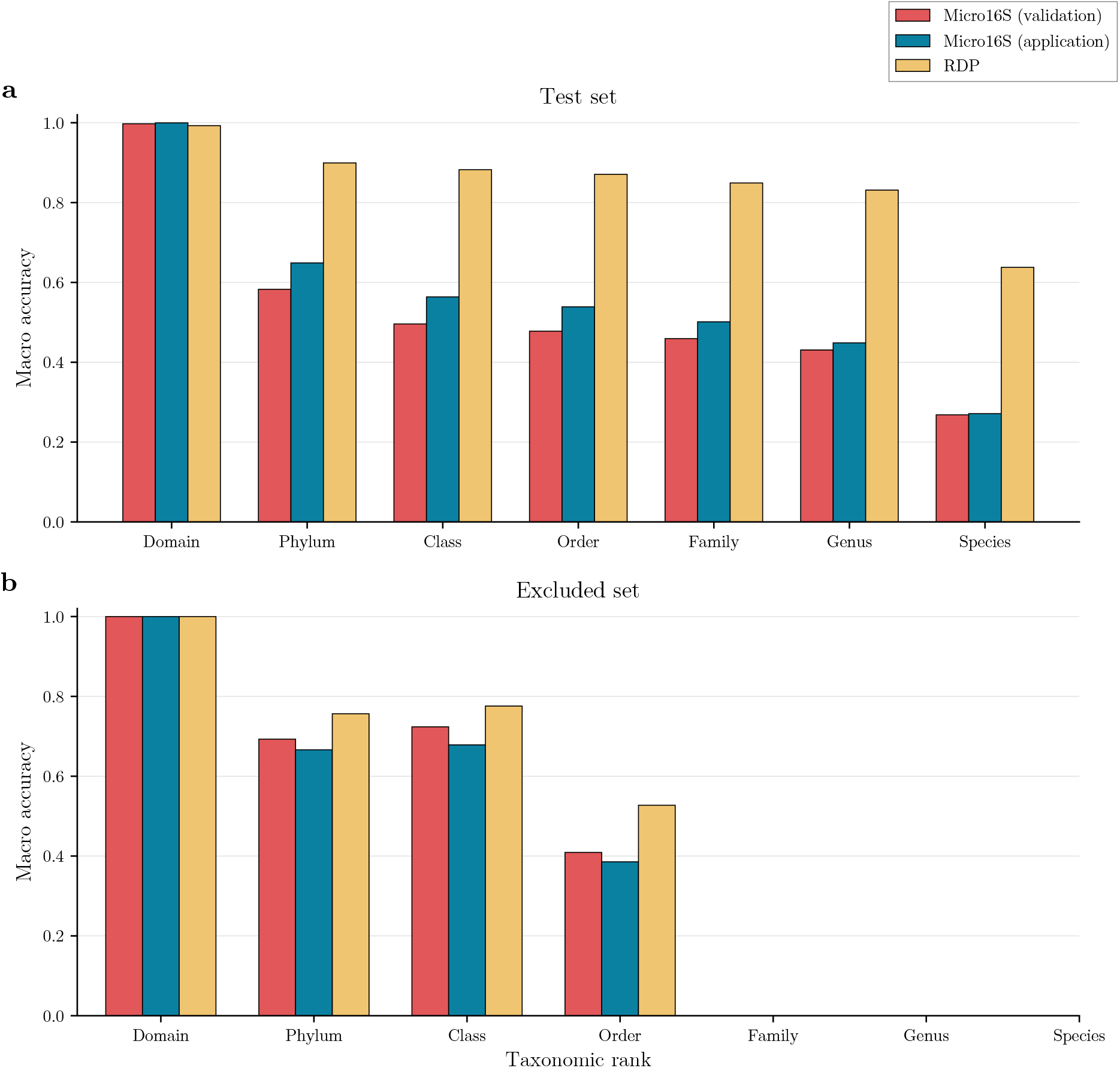
Macro accuracy scores for Micro16S and RDP across taxonomic ranks on the test and excluded sets. **a** Test set macro accuracies shown across all taxonomic ranks with red bars showing Micro16S (validation model), blue bars showing Micro16S (application model), and yellow bars showing RDP. **b** Excluded set macro accuracies using the same arrangement and colours.

**Table 1.** Clustering performance (V-measure) under mixed-region versus single-random-region sampling for the validation model on train and test sets.

| Model | Dataset split | Region sampling | Domain | Phylum | Class | Order | Family | Genus |
| --- | --- | --- | --- | --- | --- | --- | --- | --- |
| Validation | Train | Mixed regions | 0.985 | 0.709 | 0.805 | 0.874 | 0.915 | 0.930 |
| Validation | Train | Single random region | 0.985 | 0.715 | 0.817 | 0.886 | 0.934 | 0.960 |
| Validation | Test | Mixed regions | 0.975 | 0.698 | 0.793 | 0.892 | 0.942 | 0.961 |
| Validation | Test | Single random region | 0.976 | 0.704 | 0.802 | 0.899 | 0.950 | 0.974 |

**Table 2.** Consensus 16S rRNA amplicon regions derived from ASVs across the AGP and HMC datasets.

| Dataset(s) of origin | Region | Archaeal<br>start | Archaeal<br>end | Bacterial<br>start | Bacterial<br>end | Min<br>length | Median<br>length | Max<br>length |
| --- | --- | --- | --- | --- | --- | --- | --- | --- |
| Multiple HMC datasets | V1-V2 | 11 | 374 | 14 | 347 | 344 | 375 | 456 |
| Multiple HMC datasets | V1-V2 | 26 | 355 | 29 | 328 | 324 | 356 | 437 |
| Multiple HMC datasets | V1-V2 | 40 | 286 | 43 | 304 | 257 | 319 | 372 |
| One HMC dataset | V1-V2 | 189 | 339 | 50 | 312 | 189 | 320 | 373 |
| One HMC dataset | V1-V2 | 47 | 395 | 50 | 368 | 339 | 377 | 459 |
| Multiple HMC datasets | V1-V3 | 14 | 490 | 17 | 520 | 487 | 542 | 613 |
| One HMC dataset | V1-V3 | 49 | 479 | 52 | 509 | 442 | 510 | 568 |
| Multiple HMC datasets | V3 | 357 | 616 | 374 | 655 | 272 | 337 | 384 |
| One HMC dataset | V3 | 367 | 595 | 384 | 655 | 267 | 327 | 374 |
| Multiple HMC datasets | V3 | 362 | 640 | 379 | 678 | 295 | 355 | 402 |
| Multiple HMC datasets | V3 | 341 | 563 | 360 | 603 | 239 | 299 | 346 |
| Multiple HMC datasets | V3 | 338 | 606 | 356 | 643 | 280 | 343 | 390 |
| Multiple HMC datasets | V3 | 362 | 590 | 379 | 624 | 240 | 301 | 348 |
| Multiple HMC datasets | V3-V4 | 338 | 724 | 356 | 784 | 421 | 484 | 531 |
| Multiple HMC datasets | V3-V4 | 363 | 728 | 380 | 791 | 407 | 467 | 514 |
| Multiple HMC datasets | V3-V4 | 362 | 701 | 379 | 760 | 377 | 437 | 484 |
| One HMC dataset | V3-V5 | 338 | 850 | 356 | 905 | 532 | 602 | 656 |
| The AGP dataset | V4 | 471 | 594 | 531 | 654 | 136 | 184 | 231 |
| Multiple HMC datasets | V4 | 471 | 724 | 531 | 783 | 265 | 313 | 363 |
| Multiple HMC datasets | V4 | 496 | 705 | 555 | 763 | 221 | 269 | 319 |
| Multiple HMC datasets | V4 | 491 | 739 | 551 | 806 | 268 | 316 | 366 |
| Multiple HMC datasets | V4 | 471 | 699 | 531 | 758 | 240 | 288 | 338 |
| Multiple HMC datasets | V4 | 465 | 675 | 526 | 735 | 222 | 270 | 320 |
| Multiple HMC datasets | V4 | 496 | 724 | 555 | 783 | 241 | 289 | 339 |
| Multiple HMC datasets | V4-V5 | 464 | 854 | 524 | 908 | 396 | 445 | 499 |
| One HMC dataset | V4-V5 | 495 | 868 | 552 | 923 | 383 | 432 | 482 |
| Multiple HMC datasets | V5-V6 | 747 | 1006 | 806 | 1062 | 271 | 318 | 370 |
| One HMC dataset | V5-V6 | 721 | 1006 | 780 | 1062 | 297 | 344 | 396 |
| One HMC dataset | V5-V6 | 723 | 1050 | 782 | 1111 | 338 | 390 | 440 |

**Table 3.** Excluded families used for excluded split, with domains and sequence counts.

| Family | Phylum | Domain | Number of sequences |
| --- | --- | --- | --- |
| Rickettsiaceae | Pseudomonadota | Bacteria | 64 |
| Azospirillaceae | Pseudomonadota | Bacteria | 70 |
| Brevibacteriaceae | Actinomycetota | Bacteria | 72 |
| Leptotrichiaceae | Fusobacteriota | Bacteria | 73 |
| Methylophilaceae | Pseudomonadota | Bacteria | 75 |
| Bacillaceae_D | Bacillota | Bacteria | 78 |
| Sporomusaceae | Bacillota | Bacteria | 90 |
| Azotimanducaceae | Pseudomonadota | Bacteria | 93 |
| Deinococcaceae | Deinococcota | Bacteria | 93 |
| Anaerolineaceae | Chloroflexota | Bacteria | 94 |
| Opitutaceae | Verrucomicrobiota | Bacteria | 94 |
| Balneatricaceae | Pseudomonadota | Bacteria | 106 |

**Table 4.** Summary statistics of transformed RED-derived phylogenetic distances used for Micro16S training, stratified by unshared taxonomic rank.

| Rank | Min | 5th percentile | Mean | 95th percentile | Max |
| --- | --- | --- | --- | --- | --- |
| Domain | 1.3574 | 1.3574 | 1.3574 | 1.3574 | 1.3574 |
| Phylum | 0.5314 | 0.8584 | 1.0110 | 1.2134 | 1.2134 |
| Class | 0.3844 | 0.5445 | 0.6880 | 0.7803 | 1.1742 |
| Order | 0.1056 | 0.2192 | 0.3640 | 0.5015 | 0.8516 |
| Family | 0.0334 | 0.0837 | 0.1810 | 0.3440 | 0.5719 |
| Genus | 0.0007 | 0.0133 | 0.0797 | 0.1640 | 0.2896 |
| Species | <0.0001 | 0.0010 | 0.0157 | 0.0435 | 0.0912 |
| Subsequence | 0.0000 | 0.0000 | 0.0000 | 0.0000 | 0.0000 |

## References

1. Woese, C. R. & Fox, G. E. Phylogenetic structure of the prokaryotic domain: The primary kingdoms. Proceedings of the National Academy of Sciences 74, 5088–5090 (1977).

2. Janda, J. M. & Abbott, S. L. 16S rRNA gene sequencing for bacterial identification in the diagnostic laboratory: Pluses, perils, and pitfalls. Journal of Clinical Microbiology 45, 2761–2764 (2007).

3. Chakravorty, S., Helb, D., Burday, M., Connell, N. & Alland, D. A detailed analysis of 16S ribosomal RNA gene segments for the diagnosis of pathogenic bacteria. Journal of Microbiological Methods 69, 330–339 (2007).

4. Knight, R. et al. Best practices for analysing microbiomes. Nature Reviews Microbiology 16, 410–422 (2018).

5. Callahan, B. J. et al. DADA2: High-resolution sample inference from Illumina amplicon data. Nature Methods 13, 581–583 (2016).

6. Wang, Q., Garrity, G. M., Tiedje, J. M. & Cole, J. R. Naïve Bayesian classifier for rapid assignment of rRNA sequences into the New Bacterial Taxonomy. Applied and Environmental Microbiology 73, 5261–5267 (2007).

7. DeSantis, T. Z. et al. Greengenes, a chimera-checked 16S rRNA gene database and workbench compatible with ARB. Applied and Environmental Microbiology 72, 5069–5072 (2006).

8. Chuvochina, M. et al. SILVA in 2026: A global core biodata resource for rRNA within the DSMZ digital diversity. Nucleic Acids Research 54, D334–D341 (2025).

9. Hug, L. A. et al. A new view of the tree of life. Nature Microbiology 1, (2016).

10. Parks, D. H. et al. A standardized bacterial taxonomy based on genome phylogeny substantially revises the tree of life. Nature Biotechnology 36, 996–1004 (2018).

11. Parks, D. H. et al. GTDB release 10: A complete and systematic taxonomy for 715,230 bacterial and 17,245 archaeal genomes. Nucleic Acids Research 54, D743–D754 (2025).

12. Lutz, K. C. et al. A survey of statistical methods for microbiome data analysis. Frontiers in Applied Mathematics and Statistics 8, 884810 (2022).

13. Asnicar, F., Thomas, A. M., Passerini, A., Waldron, L. & Segata, N. Machine learning for microbiologists. Nature Reviews Microbiology 22, 191–205 (2023).

14. Jiang, H., Chen, Y., Hu, Y., Wang, Z. & Lu, X. Soil bacterial communities and diversity in alpine grasslands on the Tibetan Plateau based on 16S rRNA gene sequencing. Frontiers in Ecology and Evolution 9, 630722 (2021).

15. Wu, H. et al. Machine learning prediction of obesity-associated gut microbiota: Identifying Bifidobacterium pseudocatenulatum as a potential therapeutic target. Frontiers in Microbiology 15, 1488656 (2025).

16. Garach Vélez, I., Ortuño Guzmán, F. M., Rojas Ruiz, I. & Herrera Maldonado, L. J. Exploring the role of normalization and feature selection in microbiome disease classification pipelines. GigaScience 14, (2025).

17. Jasner, Y., Belogolovski, A., Ben-Itzhak, M., Koren, O. & Louzoun, Y. Microbiome preprocessing machine learning pipeline. Frontiers in Immunology 12, 677870 (2021).

18. Zhang, J. et al. Phylo-Spec: A phylogeny-fusion deep learning model advances microbiome status identification. mSystems 10, e01453–25 (2025).

19. Zhai, J. et al. DeepBiome: A phylogenetic tree informed deep neural network for microbiome data analysis. Statistics in Biosciences 17, 191–215 (2024).

20. Wang, B., Shen, Y., Fang, J., Su, X. & Xu, Z. Z. DeepPhylo: Phylogeny-aware microbial embeddings enhanced predictive accuracy in human microbiome data analysis. Advanced Science 11, (2024).

21. Woloszynek, S., Zhao, Z., Chen, J. & Rosen, G. L. 16S rRNA sequence embeddings: Meaningful numeric feature representations of nucleotide sequences that are convenient for downstream analyses. PLOS Computational Biology 15, e1006721 (2019).

22. Vaswani, A. et al. Attention is all you need. (2017) doi:10.48550/ARXIV.1706.03762.

23. Devlin, J., Chang, M.-W., Lee, K. & Toutanova, K. BERT: Pre-training of deep bidirectional transformers for language understanding. (2018) doi:10.48550/ARXIV.1810.04805.

24. Dosovitskiy, A. et al. An image is worth 16×16 words: Transformers for image recognition at scale. (2020) doi:10.48550/ARXIV.2010.11929.

25. He, K. et al. Masked autoencoders are scalable vision learners. (2021) doi:10.48550/ARXIV.2111.06377.

26. Pope, Q., Varma, R., Tataru, C., David, M. M. & Fern, X. Learning a deep language model for microbiomes: The power of large scale unlabeled microbiome data. PLOS Computational Biology 21, e1011353 (2025).

27. Nalbantoglu, O. U., Ermis, B. H. & Gundogdu, A. Deep learning-enhanced wellness scores: A population level study on gut microbiome profiling and health prediction. Biomedical Signal Processing and Control 110, 108146 (2025).

28. Zhang, H. et al. MGM as a large-scale pretrained foundation model for microbiome analyses in diverse contexts. Advanced Science https://doi.org/10.1002/advs.202513333 (2026) doi:10.1002/advs.202513333.

29. Medearis, N. A., Zhu, S. & Zomorrodi, A. R. BiomeGPT: A foundation model for the human gut microbiome. https://doi.org/10.64898/2026.01.05.697599 (2026) doi:10.64898/2026.01.05.697599.

30. Abdill, R. J. et al. Integration of 168,000 samples reveals global patterns of the human gut microbiome. Cell 188, 1100–1118.e17 (2025).

31. Schroff, F., Kalenichenko, D. & Philbin, J. FaceNet: A unified embedding for face recognition and clustering. in 2015 IEEE conference on computer vision and pattern recognition (CVPR) 815–823 (IEEE, 2015). doi:10.1109/cvpr.2015.7298682.

32. Hermans, A., Beyer, L. & Leibe, B. In defense of the triplet loss for person re-identification. (2017) doi:10.48550/ARXIV.1703.07737.

33. McDonald, D. et al. American Gut: An open platform for citizen science microbiome research. mSystems 3, e00031–18 (2018).

34. Cuesta-Zuluaga, J. de la et al. Age- and sex-dependent patterns of gut microbial diversity in human adults. mSystems 4, e00261–19 (2019).

35. Gulati, A. et al. Conformer: Convolution-augmented transformer for speech recognition. (2020) doi:10.48550/ARXIV.2005.08100.

36. Su, J. et al. RoFormer: Enhanced transformer with rotary position embedding. (2021) doi:10.48550/ARXIV.2104.09864.

37. McInnes, L., Healy, J. & Melville, J. UMAP: Uniform manifold approximation and projection for dimension reduction. (2018) doi:10.48550/ARXIV.1802.03426.

38. Cuturi, M. Sinkhorn distances: Lightspeed computation of optimal trans-portation distances. https://doi.org/10.48550/ARXIV.1306.0895 (2013) doi:10.48550/ARXIV.1306.0895.

